# Cryomilling Tethered Chromatin Conformation Capture reveal new insights into inter-chromosomal interactions

**DOI:** 10.1101/2022.02.03.478915

**Authors:** Jiang Xu, Sanjeev Kumar, Nan Hua, Yi Kou, Xiao Lei, Michael P. Rout, John D. Aitchison, Frank Alber, Lin Chen

## Abstract

Traditional methods used to map the three-dimensional organization of chromatin in-situ generally involve chromatin conformation capture by formaldehyde crosslinking, followed by detergent solubilization and enzymatic digestion of DNA. Ligation of proximal DNA fragments followed by next generation sequencing (NGS) generates contact information that enables a global view of the chromatin conformation. Here, we explore the use of cryomilling to physically fragmentize the cells under cryogenic conditions to probe chromatin interactions in the cryomilled cell fragments by the tethered chromatin conformation capture (TCC). Our results show that cryomilling TCC (CTCC) can generate a global contact map similar to that obtained with in-situ Hi-C. This result suggests that summation of chromatin interactions mapped in individual subcellular fragments can reconstitute the global contact map of intact cells in an ensemble manner, paving the way for chromatin conformation analyses of solid tissue by CTCC. Compared with the conventional in-situ methods such as Hi-C, CTCC shows more uniform access to different subcompartments of the folded genome. On the other hand, most inter-chromosomal (trans) contacts are diminished or lost in CTCC except for a group of unique trans contacts that remain intact throughout the cryomilling and in-vitro crosslinking steps. These apparently ultra-stable trans interactions have much enhanced signal in CTCC due to the elimination of signals of most, presumably weak and transient trans interactions. Systematic and comparative analyses between CTCC and in-situ Hi-C provide further insights into the chromatin structure organization and reveal a generally unentangled chromosome interface and the existence of stable inter-chromosomal contacts that may represent intermingled inter-chromosomal interfaces.

## Introduction

How chromatin organizes inside the nucleus and how such organization affects the regulation of gene expression remain important but challenging questions to address. With the advent of NGS techniques, in particular in combination with the 3C technique(1–3), the field has begun to obtain important insights into how chromatin is arranged in three dimensional space, and how chromatin structure may impact the transcriptional activity of genes. Within the last decades, many structural features of the interphase nucleus, such as chromatin compartments(2), topological associating domains (TAD)(4, 5)(4), subcompartments and loops(6), and micro-TADs(7, 8) have been identified by 3C-based technologies. These subnuclear structures, some of which have previously been observed by microscopic and cell biological studies, have also been confirmed by other genomics techniques, notably GAM(9), SPRITE(10) and ChIA-Drop(11) and imaging studies using 3D FISH augmented by super resolution microscopy techniques(12–14). However, biochemical characterization of these chromatin structural features, such as the physical stability of chromatin contacts belonging to different subcompartments, has been generally lacking. Moreover, discrepancies became evident when comparing the results of 3C-based techniques with those from imaging analyses(5, 15), raising the question that key steps of the 3C-based techniques, including formaldehyde crosslinking, SDS treatment and nuclease digestion, may have inherent limitation in chromatin conformation analysis.

In order to address these issues, many variants of the original protocol have been attempted with or without formaldehyde crosslinking, and with or without SDS treatment(6),(16),(17),(18). Indeed, it has been shown that nuclease digestion has uneven accessibility and/or efficiency across different regions of the nucleus(19). One potential solution to this problem is to physically open up the nucleus to allow more even sampling of the genome by various mapping methods. Sonication is currently one of the most widely adopted cell fragmentation methods in various chromatin related approaches, such as ChIP-Seq(20), ChIA-PET(21), Hi-ChIP(22), PLAC-seq(23), SPRITE(10), ChIA-Drop(11). However, high shearing forces and heating could potentially denature nuclear proteins and destabilize chromatin assemblies, which could distort conformational analysis. In addition to the nuclease accessibility issue, chromatin conformation capture in intact cells by formaldehyde crosslinking is a poorly understood process that could introduce bias and artifacts in 3C-based protocols(18, 24).

Formaldehyde cross-linking is a slow and difficult-to-control process, requiring tens of minutes to generate stably crosslinked chromatins. Its diffusion and crosslinking efficiency could vary in different nuclear regions. Thus, nearly two decades after the introduction of the 3C method and subsequent development and wide-spread adoption of the Hi-C or Hi-C-like approaches, a major challenge in the field of chromatin conformation study remains finding methods that can faithfully capture native nuclear interactions that drive the assembly and function of the nuclear structure.

Here, we present a new technique, called CTCC, which combines cryomilling with tethered chromatin conformation capture (TCC), a 3C-based technique we previously developed to probe the structure of the chromatin(3). Cryomilling is a mechanical breaking method that has long been used in molecular biological studies(25–29), in which cells are flash frozen and mechanically ground into fine powder under ultra-low temperature (e.g. in liquid nitrogen or argon). With flash freezing, cellular structures including chromatin conformations can be captured instantaneously. With cryomilling, cells are fragmentized into nanometer to sub-micrometer particles in the frozen state to maximally preserve native structures and interactions in each cell fragment. These cell fragments could be analyzed directly for DNA content using the barcoding strategy such as in SPRITE(10) or protein composition and structure using antibody enrichment followed by mass spectrometry(30). In this study, we will focus on DNA proximity mapping using 3C-based methods in the cryomilled cell fragments, wherein the access to chromatin complexes is much enhanced by the physical breaking and opening of the cellular structure. Different from other in-vitro techniques, such as GAM(9), SPRITE(10) and CHIA-Drop(11), CTCC combines physical breaking and proximity ligation, thereby measuring DNA-DNA proximity directly in fragmentized subcellular fragments rather than relying on the inference of co-localization of DNA regions on the same particle or slice. In this study, we show that CTCC can be an effective chromatin conformation mapping technique that complements existing mapping techniques. Through systematic and comparative analyses between CTCC and Hi-C or between CTCC experiments with different experimental settings, our studies also shed new insights into the structure and function of chromatin organization in the nucleus.

## Results

### Cell and nuclear fragmentation by cryomilling

The basic idea behind CTCC is to capture the cellular structure by flash freezing and subsequently break down the frozen cells under cryogenic conditions (**Figure 1a**). The resulting fragments, including the nuclear fragments, can be analyzed either by structural techniques (e.g., cryo-electron microscopy or cryo-electron tomography), or by molecular methods after chemical stabilization upon thawing. In principle, cryomilling can fragmentize cells into subcellular particles of various sizes by controlling the cryomilling conditions. In the current study, we used an extended cryomilling time of 86 minutes to break up cells into fine particles. A key issue encountered in developing the CTCC protocol is the high aggregation tendency of the cryomilled cell fragments, which behaved like colloidal particles when mixed with buffer solution and formed conglomerates of various sizes. This could be a serious problem in chromatin contact mapping as ligation between DNA from different particles could lead to false proximity information. Two approaches were used to overcome this hurdle. One is to use SDS to prevent the aggregation of cell fragments. In the cases wherein SDS was not used, a buffer with an optimized pH and ionic strength was used to establish a sheath of charge around the cryomilled cell particles so that they stay as highly dispersed fine particles in solution (**Supplementary Video**). Once stabilized, the cryomilled subcellular fragments can be analyzed and subject to chromatin contact mapping.

**Figure 1.**
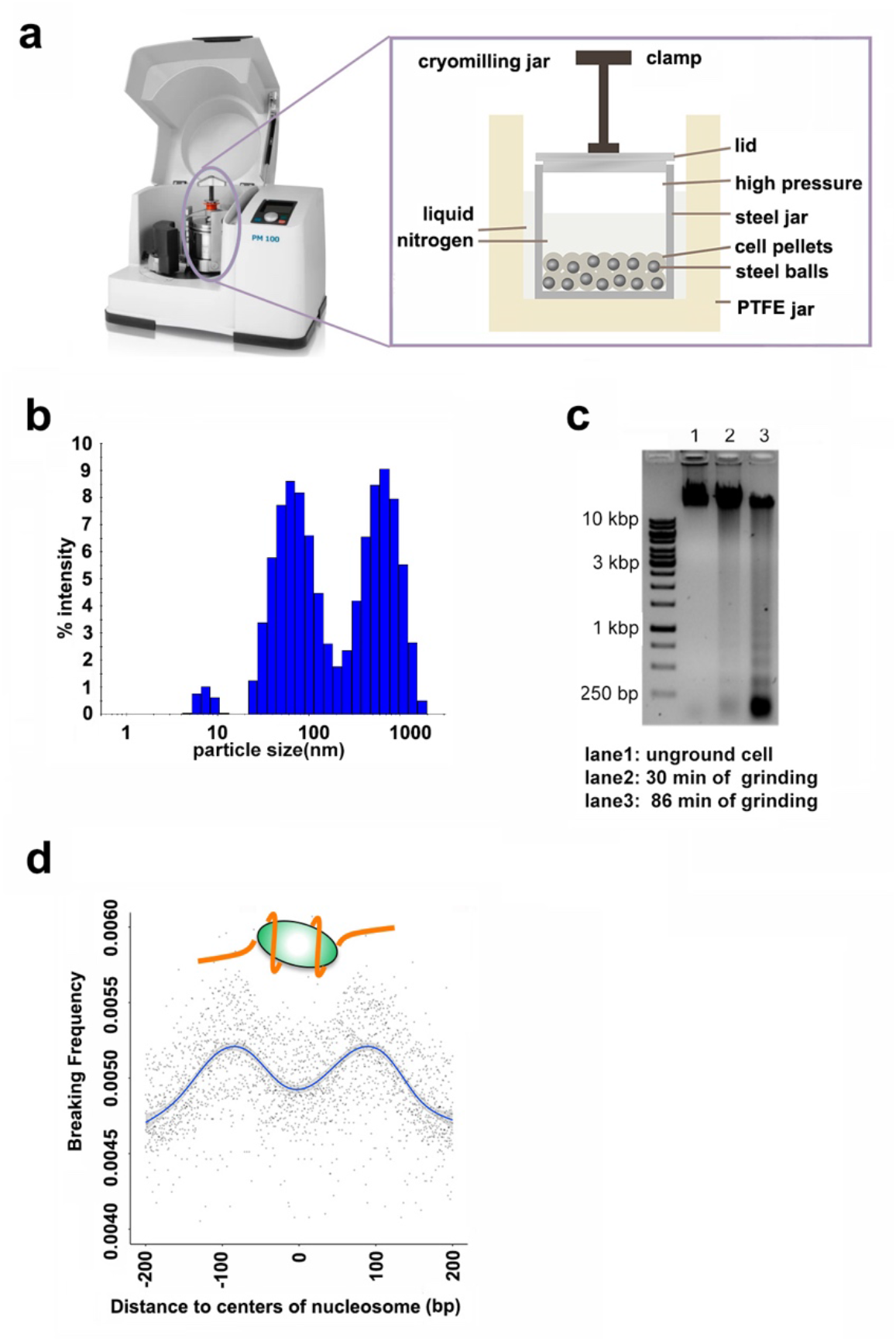
**a**. Cryomilling machine setup: a commercial milling machine (Retsch Tech PM-100) was equipped with a custom crafted cryomilling jar that enables long time cryomilling. **b**. Dynamic light scattering (DLS) measurement of the size distribution of formaldehyde solution fixed, cryomilled GM12878 cells with 86 min cryomilling protocol. **c**. Agarose gel electrophoresis of purified genomic DNA from cryomilled GM12878 cell power. **d**. Pileup analysis of DNA breaking frequencies relative to the center positions of nucleosomes

Analyses by dynamic light scattering (DLS) revealed two dominant populations centered around 60 nm and 400 nm, respectively (**Figure 1b**). This two-peak distribution of particle size is surprising given the expected random fragmentation of the cells by cryomilling. One possible explanation is that the two distinct particle size populations may correspond to two different subcellular regions (e.g., cytoplasmic, and nuclear) that have different cryomilling efficiency. Given the extended cryomilling time used, this size range may represent the lower limit of the current cryomilling instrument used in our studies. Purified genomic DNA from the cryomilled cell particles showed DNA fragment sizes in the range of ∼200 bp and its multiples (**Figure 1c**). pileup analysis using reference nucleosome positions(31) showed that cryomilling preferably breaks the DNA on the edge of nucleosomes (**Figure 1d**), which explains the ladder pattern observed by DNA gel electrophoresis. Comparison of cryomilling with sonication and micrococcal nuclease (MNase) digestion reveals different DNA breaking positions relative to the binding sites of nucleosomes (**Supplementary Figure 1a**) and transcription factors (**Supplementary Figure 1b**), indicating different DNA breakage preferences between these methods. Overall, these experiments show that cryomilling could efficiently break down the cells as judged by the particle size and DNA fragment length.

### Adapting cryomilling to tethered chromatin conformation capture

To facilitate the mapping of chromatin contacts in the subcellular particles generated by cryomilling, the general framework of TCC was adopted by tethering the subcellular particles on carrier beads to perform proximity ligations. Because of its in-vitro and modular nature, parameters of the CTCC protocol, including the cryomilling time, detergent usage, crosslinking chemicals, and conditions, etc., could be adjusted according to various experimental considerations (see below) (**Figure 2a**). For example, in one representative CTCC experiment, 1.44 billion (out of 1.8 liters of cell culture) of freshly grown GM12878 cells were resuspended in isotonic glucose solution to a 50% slurry, and flash frozen drop by drop in liquid nitrogen and cryomilled for 86 min. The cryomilled cell powder was added to a crosslinking buffer containing 1% formaldehyde at room temperature for 15 min. The crosslinked cell fragments were then subject to HindIII digestion, end labelling and proximity ligation (See methods for more details). Sequencing of the resulting library of chimeric DNA fragments reveals a contact frequency map (**Figure 2b**) and mapping statistics similar to that obtained with traditional Hi-C (**Supplementary Figure 2**). CTCC detected all known features seen with traditional dilution Hi-C and in-situ Hi-C, such as TAD, subcompartments, strips and loops (**Figure 2b**). These experiments demonstrate that chromatin contacts mapped in randomly generated individual subcellular fragments, when summed together, could recapitulate the ensemble global chromatin conformation features observed with Hi-C on populations of cells.

**Figure 2.**
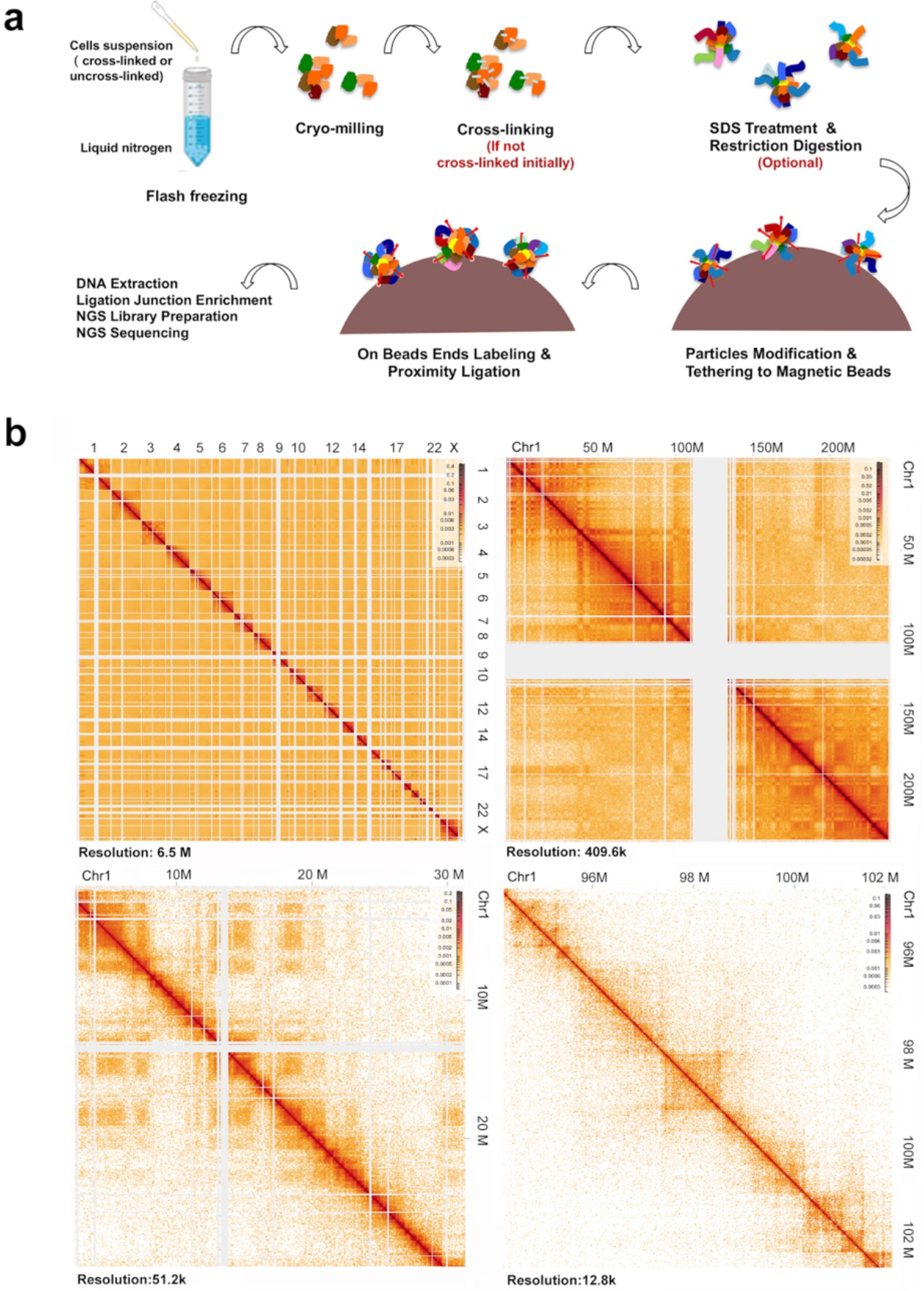
**a**. Flow chart illustration of CTCC experiments. **b**. Contact maps of CTCC (with HindIII) at different resolutions show typical known 3C features. (Upper-left: overall, upper-right: compartment, lower-left: Topological Associated Domains (TADs), lower-right: loop and strip.

### Enhanced access and sampling of the genome by CTCC

CTCC showed enhanced access to the genome as a result of the physical breaking of the cells as compared with conventional dilution Hi-C or in-situ Hi-C. This is not only evident from the evenly distributed DNA breaks generated by cryomilling (**Figure 3a**, top track) but also from the DNA breaks generated by HindIII digestion of cryomilled cell fragments (**Figure 3a**, middle track, cryomilling aided HindIII digestion) as compared with those from in-situ digestion of the genomic DNA (**Figure 3a**, bottom track, HindIII alone). DNA digestion by restriction enzymes (e.g., HindIII and MobI) are generally less efficient in some subcompartments (e.g., B1, B2 and B3) than in others (e.g., A1 and A2). However, these differences are substantially reduced by cryomilling. (**Supplementary Figure 3a**). By plotting DNA break frequencies in terms of breaking points per kilobase per million mapped reads (BPKM), it becomes clear that cryomilling aided HindIII (**Figure 3b, right panel**) not only generated more breaking points but also a more narrowly distributed density than in-situ HindIII digestion alone (**Figure 3b, left panel**). The differences are more prominent in the B subcompartments (B1, B2 and B3) as compared with the A (A1 and A2) compartments. These analyses suggest that CTCC samples the genome more efficiently and evenly across different subcompartments than in-situ Hi-C. The division matrix between CTCC and dilution Hi-C(32) also shows that dHi-C has more signal in compartment A of the genome, while CTCC enhances signal in compartment B of the genome (**Figure 3c, Supplementary Figure 3b**). These observations are consistent with the idea that A subcompartments (A1, A2) generally represent open and more accessible chromatin regions whereas B subcompartments (B1, B2 and B3) are more compact and less accessible chromatin regions. Although overall loops and TAD boundaries are similar between Hi-C and CTCC (**supplementary Figure 3c**), pileup analyses of loops and TAD boundaries from different subcompartments show that the signals for the closed subcompartments in CTCC substantially increased as compared with Hi-C (**Figure 3d**). Together, these results show that cryomilling facilitates the probing of chromatin conformation features in the closed subcompartments of the genome.

**Figure 3.**
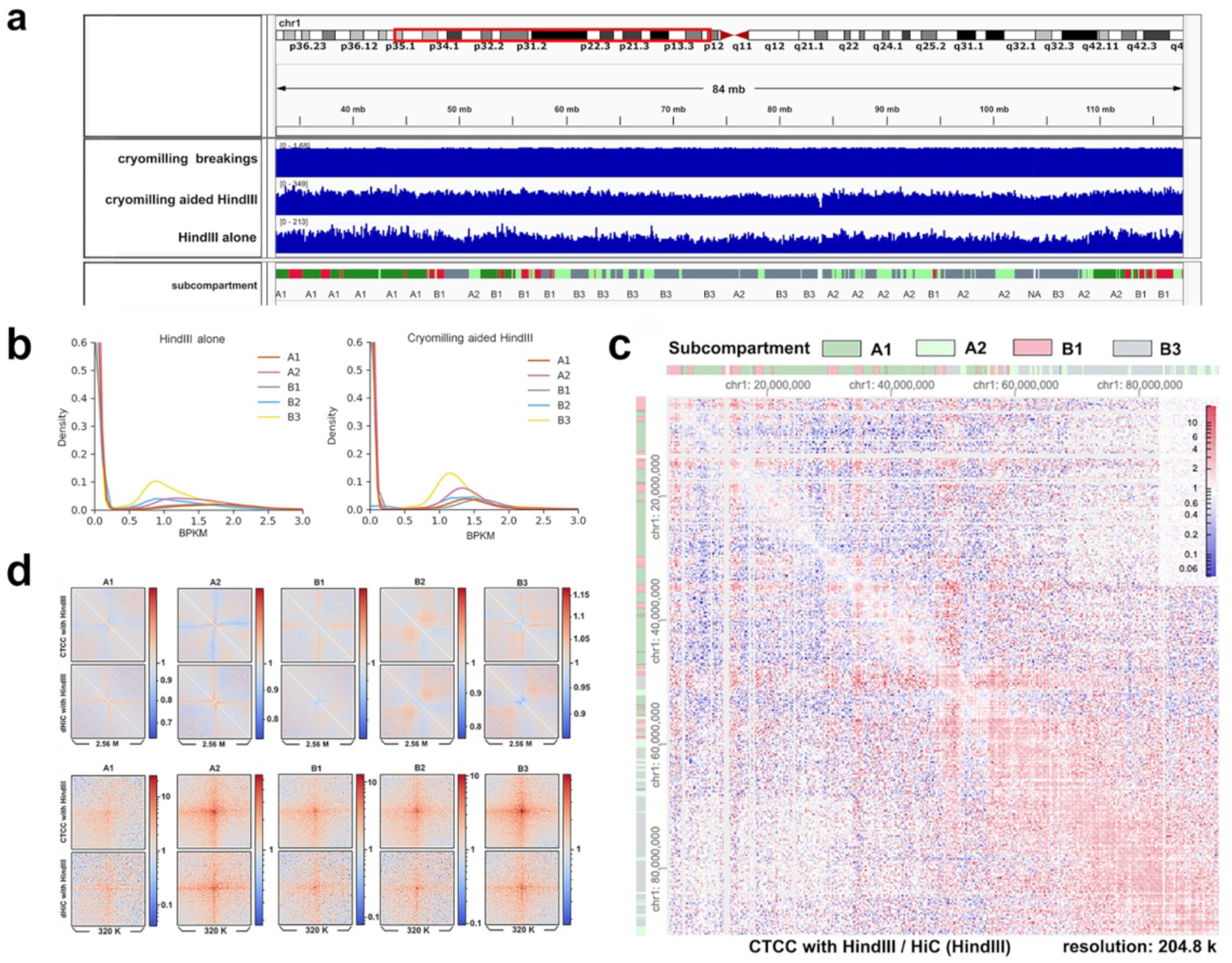
**a**. Comparison of DNA breaks features between cryomilling alone, cryomilling aided HindIII digestion and HindIII digestion alone across different subcompartment of chromosome 1 (viewed in IGV). **b**. Kernel density estimate plot (kde-plot) comparison of DNA breaking frequency of consecutive 1 kb bins across the genome between cryomilling aided HindIII digestion and HindIII digestion alone. **c**. Division of the contact matrix of CTCC with HindIII with that of HiC with HindIII, more cis signal observed between loci within B1 and B3 subcompartments for CTCC and more signal between loci within A1 subcomparments for dHi-C **d**. Comparison of pileups of contact matrices along the diagonal at l local insulation valleys (upper panel), and loops (lower panel) between CTCC (with HindIII) and Hi-C (with HindIII).

### Comparative analyses of different subtypes of chromatin interactions mapped by CTCC and Hi-C

A major difference between CTCC and Hi-C is that CTCC is an in-vitro approach while Hi-C is an in-situ approach. Because of the in-situ nature of Hi-C, loci that are not physically associated but are close in space (**Supplementary Figure 4a, loci A and B**) could still be captured in Hi-C, while CTCC has the potential to enrich physical contact (**Supplementary Figure 4a, loci C and D**). However, in practice, we found decreased contrast in the CTCC contact map compared with Hi-C, which could be a result of interparticle ligation events in CTCC, as shown by the analysis result below.

We compared the overall chromatin contact pattern between CTCC (with cryomilling) and Hi-C (without cryomilling). CTCC As shown in the division matrix of CTCC over Hi-C (**Figure 4a**), the intra-chromosomal (cis) contacts (red boxes along the diagonal line) were well captured and apparently enhanced by CTCC. On the other hand, the inter-chromosomal (trans) contacts, corresponding to the off-diagonal blue regions, were substantially diminished in CTCC. However, some of the off-diagonal regions did show increased signal in CTCC, as indicated by red pixel (ratio of CTCC signal/Hi-C signal > 1). Whether these enhanced off-diagonal signals represent true stable trans contacts or result from increased background noises is a key question in evaluating the performance of CTCC. To address this question, we take advantage of established chromatin organization features previously observed in GM12878 to evaluate these enhanced trans interactions. Generally, a chromatin region tends to contact other intra-chromosomal (cis) or inter-chromosomal (trans) regions of the same subcompartment. Higher cross subcompartment ligation events could indicate a higher background noise in the mapping experiment. Taking the p-arm of chromosome 1 for example, which consists of mostly actively transcribed A1 subcompartment, the division matrix of CTCC/Hi-C showed that the p-arm of chromosome 1 has many trans contacts that appears to be enhanced by CTCC (the extended red streak off diagonal of the p-arm of chromosome 1 in **Figure 4a**). However, most of these trans-contacts are between different subcompartments (referred to as cross-subcompartment contacts hereafter) (**Figure 4b**, zoom in of the boxed region in **Figure 4a**), suggesting that these trans-contact signals elevated in CTCC are likely noise, such as those associated with ligation events between different particles.

**Figure 4.**
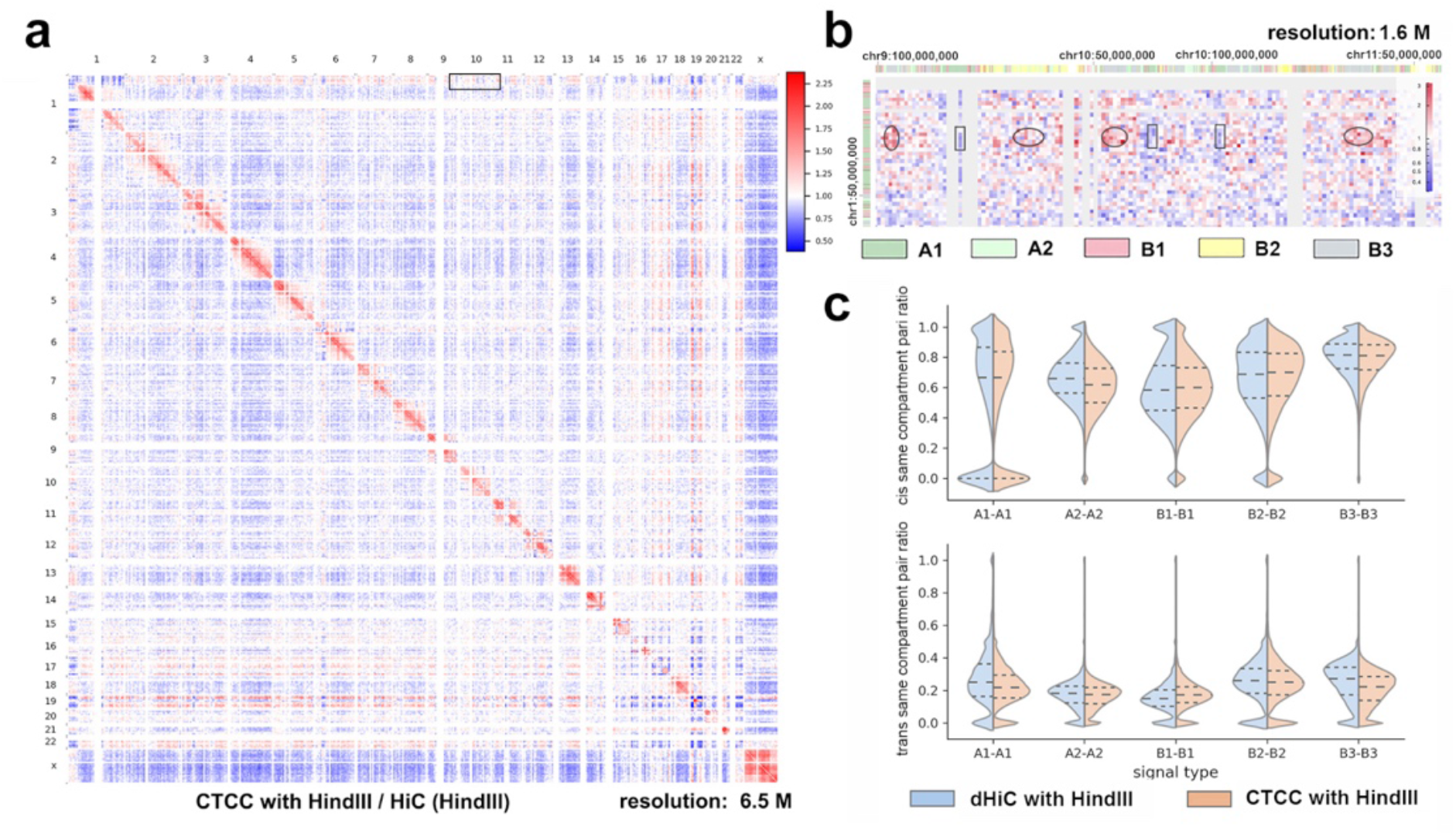
Cryomilling’s effects on signal and noise. **a**. Division of the contact matrix of CTCC with HindIII with that of Hi-C with HindIII at 6.5 M resolution across the whole genome. **b**. Zoom-in of a small area (enclosed in a small black rectangle box in a, upper middle), trans contacts between loci of A1 subcompartment are enclosed with black rectangle, trans contacts between loci of A1 subcompartment and other subcomparments are enclosed with black ellipse). **c**. Violin graph comparison of dilution HiC and CTCC (both with HindIII) in terms of “bin’s cis same compartment pair ratio” and “ bin’s trans same compartment pair ratio”, showing the distribution of the number of bins with the same “same compartment pair ratio” (see methods)

To further analyze such cross-subcompartment contacts, we calculate for a given bin (i.e., chromosome region) the ratio of its contact pairs to regions of the same subcompartment to its total contact pairs, which is defined as the bin’s “same compartment pair ratio” (see method). To evaluate the impact of cross-subcompartment ligation in intra- and inter-chromosomal contact mapping, the bin’s same compartment pair ratio can be further divided into bin’s cis same compartment pair ratio and bin’s trans same compartment pair ratio. As shown in Figure 4c, the number of bins with high “same compartment contact ratio” was substantially decreased by cryomilling. The median and quartile values of the bins cis- or trans-same compartment pair ratio were decreased by cryomilling for most subcompartments, indicating a generally higher cross-subcompartment noise in CTCC as compared with dilution Hi-C. The most dramatic decreases were seen in the A1 and A2 subcompartments (for both the cis- and trans-contacts) and in the trans-contacts of the B3 compartment, suggesting that mapping of chromatin interactions in these regions by CTCC are particularly prone to the cross-subcompartment noises mentioned above. By contrast, the B subcompartments, B1 and B2 (both the cis- and trans-contacts) and the cis- contacts of the B3 compartment, which account for most of the contacts in B3, do not show significant decrease of bins of “high same compartment contacts ratio”. Interestingly, the median and quartile values of the bins cis ‘same compartment contact ratio’ for the B1 and B2 subcompartment even showed slight increase in CTCC, this is probably because cryomilling increases the accessibility of these closed subcompartments thereby enhancing the mapping of cis contacts within. Surprisingly, the median and quartile values of the bins trans same compartment contacts ratios for the B1 subcompartment were increased substantially by cryomilling. Even though these trans contacts account for only a minor fraction of the B1 subcompartment contacts, it does raise the intriguing question if these trans contacts of B1 subcompartments represent stable inter-chromosomal interactions that are resistant to cryomilling and enhanced by CTCC.

Given the fact that trans contacts seem to be more prone to background noises associated with CTCC, we further evaluated the signal to noise ratio between CTCC and dHi-C by calculating trans pairs composition in each subcompartment (see method) (**Supplemented Figure 4b**). For a given subcompartment, all trans contacts within the same subcompartment normalized against the total trans contacts can be taken as the signal, whereas all trans contacts to other subcompartments normalized against the total trans contacts can be taken as the noise. These analyses show that the ‘signal to noise ratio’ for the B3 and A1 subcompartments was substantially decreased by cryomilling (**Supplemented Figure 4b**, left column). Further analysis shows that the noise fraction is increased more than the signal fraction for A1 subcompartment by cryomilling, causing the signal to noise ratio to decrease (**Supplemented Figure 4b**, compare the middle and right column). On the other hand, even though cryomilling decreases the noise pair fraction for the B3 subcompartment (**Supplemented Figure 4b**, right column), the signal pair fraction for the B3 subcompartment drops even more (**Supplemented Figure 4b**, middle column), causing significant drop of signal to noise ratio for the B3 (**Supplemented Figure 4b**, left column).

The above analyses suggest that compared with Hi-C, CTCC enhances the mapping of cis contacts but leads to generally reduced trans signal and more noise in the contact map. While the active subcompartments A (A1 and A2) are more prone to the cross-subcompartment noises caused by cryomilling than the inactive B subcompartments (B1, B2 and B3 cis contacts) (**Figure 4c**, upper panel), the dramatically decreased trans signal to noise ratio in terms of bin’s same compartment trans pair ratio (**Figure 4c**, lower panel) and trans pairs’ signal to noisel ratio (**Supplementary Figure 4b**) of B3 subcompartment suggest that inter-chromosomal interactions formed by the B3 subcompartment may not be stable in the cryomilling process and associated in-vitro proximity ligation steps of CTCC. On the other hand, a surprising result from these analyses is the apparent enhancement of trans contacts in the B1 subcompartment (**Figure 4c**, lower panel, Supplementary Figure 4a), suggesting the existence of stable inter-chromosomal contacts that are cryomilling resistant (discussed further below).

### Assess chromatin interaction capture by formaldehyde crosslinking

The comparative analyses between CTCC and Hi-C above suggest that CTCC carried out under different biochemical conditions (e.g., crosslinking, restriction digestion, SDS treatment and RNase digestion etc.) could provide unique opportunities to characterize different subtypes of chromatin interactions and their roles in chromatin organization. With CTCC, chromatin crosslinking could be performed either on intact cells in-situ before cryomilling or on cryomilled subcellular particles in-vitro after cryomilling. To see how chromatin interactions could be affected by the two different modes of crosslinking, we carried out parallel CTCC experiments with formaldehyde crosslinking performed before (in-situ crosslinking CTCC) or after (in-vitro crosslinking CTCC) cryomilling. Comparison of the results from these two experiments showed no significant difference in the calculated insulation scores (**Supplementary 5a**, left panel, Pearson correlation test). Pileup analyses of the local feature of insulation score valleys are also similar (**Supplementary 5a**, right panel). These results indicate that gross, large-scale genomic structural features are preserved in the cryomilling process and can be stabilized by in-vitro crosslinking of the subcellular fragments. However, the contact frequency vs genomic distance curves between the two protocols are very different, with the in-vitro crosslinking CTCC showing substantially reduced short range contacts between 1 kb to 20 kb than the in-situ crosslinking CTCC (**Figure 5a**). This difference is consistent across all subcompartments and seems more significant for subcompartment A1 and B1 (**Supplementary 5b**). Loops from either division matrix (**Figure 5b**, left) or pileup analyses (**Figure 5b**, right) also show reduced signals in the in-vitro crosslinking CTCC than the in-situ crosslinking CTCC. These observations suggest that a large amount of short to medium range interactions (1-20 kb) are disrupted by the cryomilling process and/or inefficiently captured by the in-vitro crosslinking step, which could also lead to the increased self-ligation (less than 1kb distance) observed in in-vitro crosslinking CTCC across all subcompartments (**Figure 5a, Supplementary 5b**).

**Figure 5.**
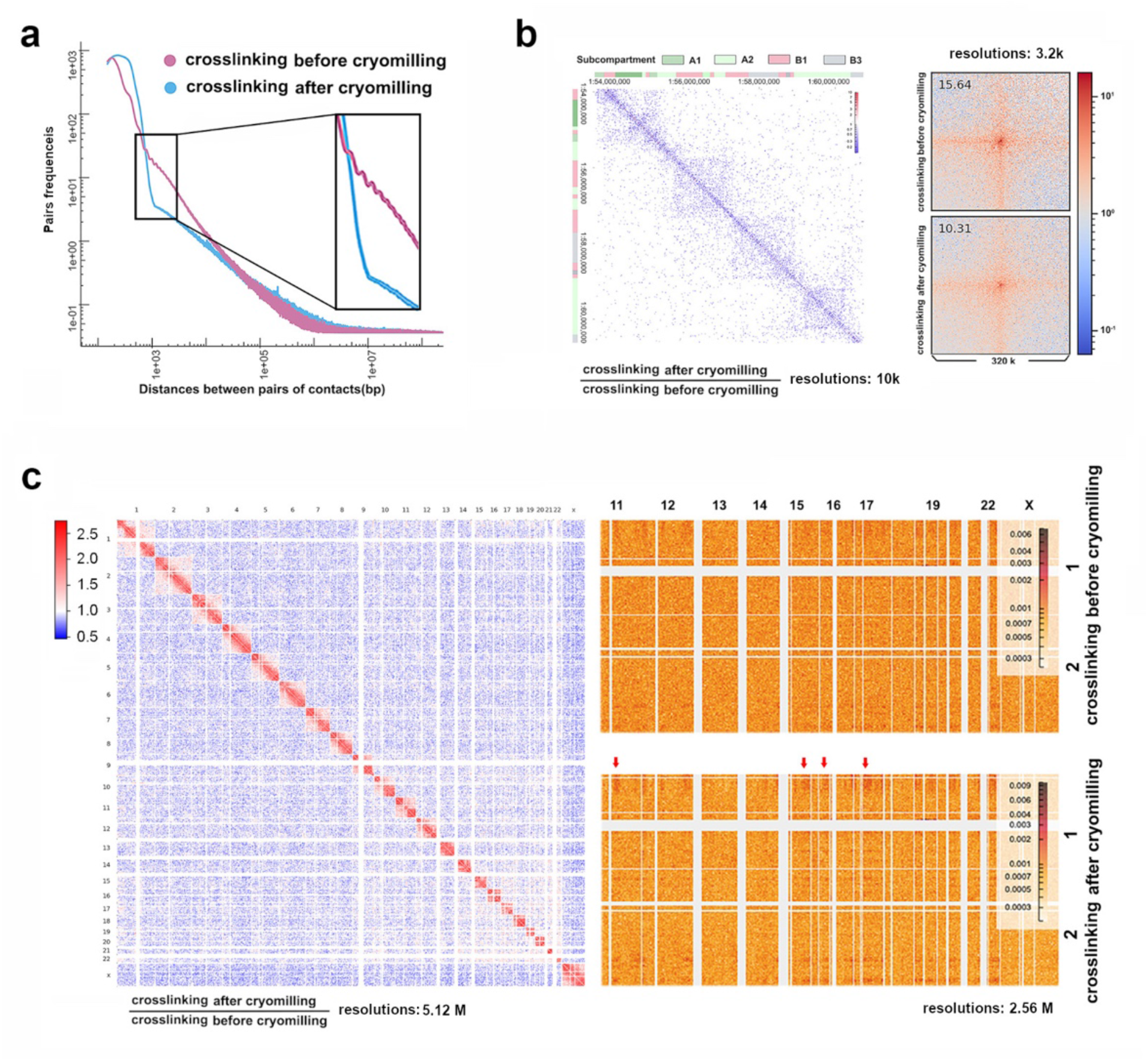
Comparison of two formaldehyde crosslinking strategies used in CTCC experiments: crosslinking before cryomilling (crosslinking in-situ) or crosslinking after cryomilling (crosslinking in-vitro) **a**. Pair-frequency-pair-distance curve comparison. **b**. Division of matrices at 10k resolution (left) and pileup of the matrices at reference loop positions, pileups use 3.2 k because the result was most eligible at this resolution. **c**. Division matrix at 5.12 M, showing overall decreased trans contacts and increased cis contacts for crosslinking after cryomilling CTCC experiment (left), while certain trans contacts stand out of the overall decreased trans contacts for crosslinking after cryomilling CTCC experiment (right, lower panel, red arrows).

Another significant difference is the presence and absence of the zig-zag feature in the contact frequency/genomic distance curve around 0.4∼2 kb range. The zig-zag pattern is prominent in the in-situ crosslinking CTCC experiment but completely absent in the in-vitro crosslinking CTCC experiment (**Figure 5a, Supplementary 5b**). The zig-zag pattern represents the short-range contacts between the neighboring nucleosomes, which was observed in previous research using in vivo approaches with formaldehyde fixed cells(7,33,34). Although the details of the interactions underlying these short-range inter-nucleosome contacts remain to be fully characterized, our analyses suggest that these interactions could be captured and stabilized by formaldehyde crosslinking in intact cells (i.e., in-situ crosslinking), probably through protein-protein crosslinking of neighboring nucleosomes. These interactions and short-range contact structures are unstable in the cryomilling process and/or inefficiently captured by the subsequent in-vitro crosslinking reaction in-vitro.

Moreover, the in-vitro crosslinking (crosslinking after cryomilling) CTCC captured much less inter-chromosomal (trans) contacts as compared with the in-situ crosslinking (crosslinking before cryomilling) CTCC, as indicated either by the much-decreased trans pair fraction in the total mapped contact pairs (**Supplementary 5c**), or by the much-reduced signal in the off-diagonal region of the contact map (**Figure 5c**). This is similar to the comparison between CTCC and Hi-C discussed above but here the difference is directly linked to different formaldehyde crosslinking protocols within CTCC. This observation suggests that most particles containing DNA from different chromosomes (hence trans contacts) tend to fall apart either in the cryomilling of non-crosslinked cells, or during the in-vitro crosslinking process. However, while most trans contacts are unstable hence unobservable in the in-vitro crosslinking CTCC, trans contacts between certain loci of the genome are surprisingly stable and become much more standing out from the overall reduced trans signals (**Figure 5c,** right, lower panel, red arrows). These stable inter-chromosomal trans-contacts may play important roles in cell functions and in organizing the inter chromosome network(35, 36), and will be further discussed below.

Overall, the above comparative analyses suggest that a large amount of short to medium range chromatin contacts and most trans interactions observed in in-situ Hi-C require formaldehyde crosslinking in the intact cells, while most of the cis chromatin interactions and some ultra-stable trans interactions remain stable in the cryomilling process and can be efficiently captured in the subsequent in-vitro crosslinking of the subcellular fragments.

### Restriction digestion of cryomillied subcellular fragments

In CTCC, DNA proximity can be mapped by ligation of DNA breaks generated by cryomilling. However, additional DNA breaks could be generated by restriction digestion of DNA in the cryomilled subcellular fragments. As shown in **Figure 6a**, restriction enzyme HindIII digestion in addition to cryomilling dramatically increases the cis contact pair frequency with distance over 1 kb. Furthermore, the enhanced mapping occurs mostly between loci of the same genomic subcompartments (**Supplementary Figure 6a**). Interestingly, different subcompartments showed different signal enhancement, with A2 and B3 subcompartments showing the most increase and A1 subcompartment showing the least increase. This observation could be explained by the mechanistic model depicted in **Supplementary Figure 6b**. Cryomilling-generated cell particles should have DNA breakings distributed on their surface, which will lead to some “intra-particle” ligation events with cryomilling alone. Restriction enzyme digestion, however, could create DNA breaks both within and on the surface of the particles, enabling more proximity ligation events within individual particles. Pileup analyses for local features around insulation score valleys, on the other hand, do not show significant difference (**Figure 6b**), indicating that this structural feature is independent of the additional restriction fragments generated by enzymatic digestion of the cryomilled subcellular particles. Loops, on the other hand, are enhanced by additional HindIII digestion (**Figure 6b**), suggesting solid physical contacts with high DNA contents, in which HindIII digestion could produce more fragments within the individual particles, leading to more ligation events around the loop structure.

**Figure 6.**
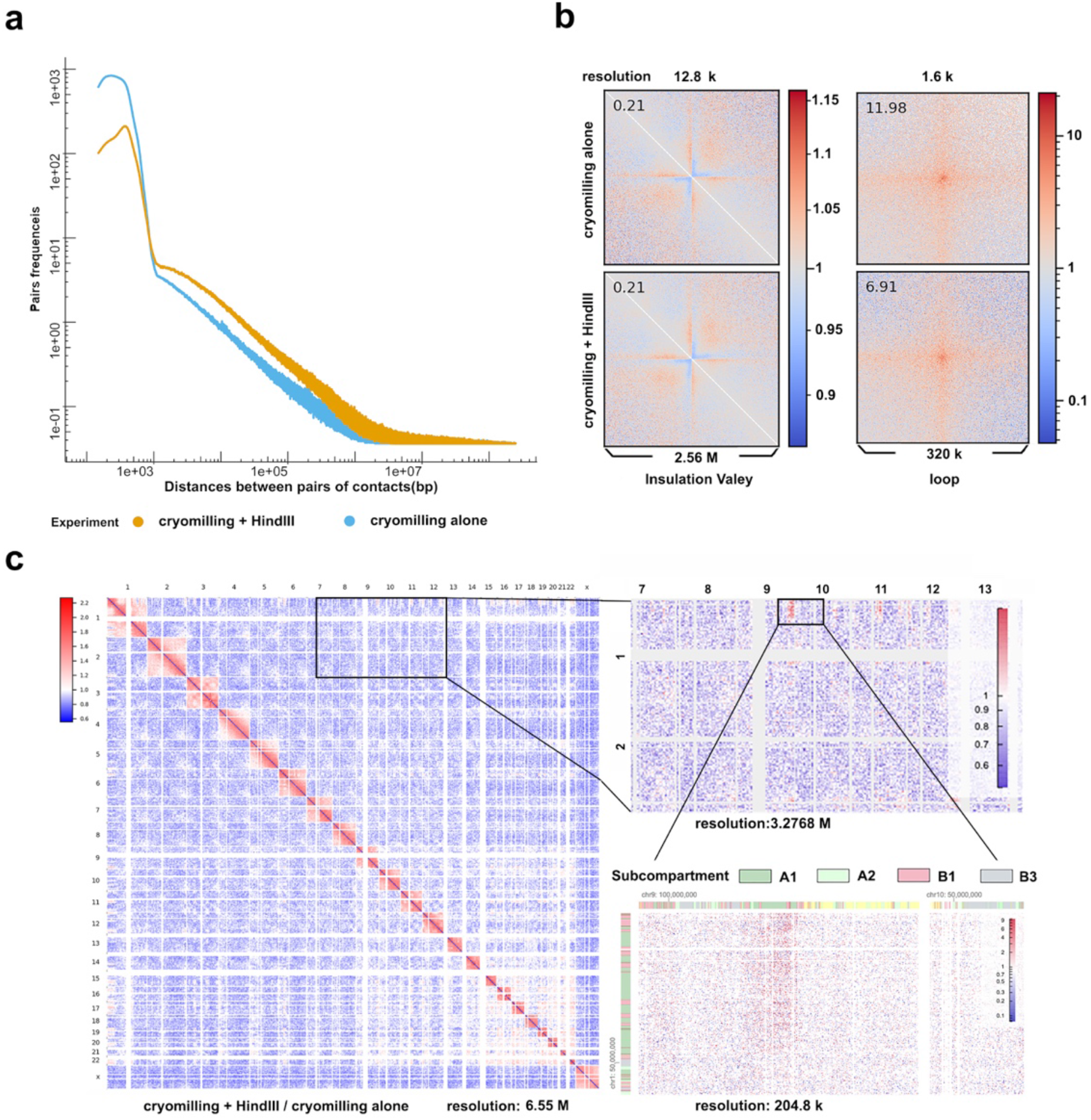
Comparison of CTCC experiments with or without HindIII digestion. **a.** Comparison of pair-frequency-pair-distance curve. **b**. Comparison of pileups of contact matrices at local insulation score valleys and reference loop positions. **c**. Division matrices between CTCC with HindIII digestion and CTCC without HindIII digestion at low and medium resolutions.

Although CTCC with or without HindIII showed overall substantially decreased trans signals (**Figure 6c**), as discussed earlier, certain trans-signals appear to be stable in the cryomilling and in-vitro crosslinking process and become standout due to the diminished overall trans signal. These cryomilling-resistant trans signals are further enhanced by HindIII digestion. The signal enhancement of these ultra-stable trans signal by Hind III digestion is similar to the those of cis signals (**Figure 6c**). This observation suggests that some of the cryomilled cell particles are composed of mostly trans interactions whose mapping could be enhanced by internal DNA digestion by restriction enzymes penetrating the particles. By calculation of pixels’ subcompartment enrichment curve (see methods, **Supplementary Figure 6c**, upper panel), we were able to show that many of the trans contacts enhanced by cryomilling/HindIII digestion belong to A1, B1 and B2, particularly the A1 subcompartments(**Supplementary Figure 6c**, lower panel). These analyses suggest that trans contacts within the A1 (or B1, B2) subcompartment are very stable and become enriched by the cryomilling and in-vitro crosslinking process of CTCC. Finally, Division of the matrix of “crosslinking after cryomilling with HindIII digestion” by those of “crosslinking before cryomilling without HindIII digestion” shows even higher overall cis value and much lower trans value, with much higher values for certain trans pixels, hence higher contrast seen on the division matrix (**Supplementary Figure 6d**, upper panel) and more significant separation of pixels’ trans subcompartment enrichment curve between A1, B1, B2 and A2, B3 subcompartments (**Supplementary Figure 6d**, lower panel). Thus, by comparing different matrix division combinations, it becomes apparent that ‘crosslinking after cryomilling’ and ‘HindIII digestion’ synergistically enhance those ultra-stable trans signal that belong to the A1, B1 and B2 subcompartments. The underlying mechanism could be explained in Supplementary Figure 6e. Because only stable contacts that can survive extensive cryomilling, remain together in the in vivo crosslinking could be further enhanced by restriction enzyme digestion, while those who are not stable could not (**Supplementary Figure 6e**). With crosslinking in-vitro (crosslinking after cryomilling) protocol, contacts, both stable and not stable, could be crosslinked by formaldehyde and survived in the solution (**Supplementary Figure 6e**). These observations thus suggest that genomic loci from A1, B1 and B2 subcompartments may form extensive stable inter-chromosomal (trans) interactions that are resistant to cryomilling and become enhanced by CTCC with in-vitro crosslinking and restriction digestion where the background associated mostly with weak trans contacts (or noise from inter-particle ligations) is diminished. Taken together, the above analyses suggest that further restriction digestion of cryomilled subcellular fragments enhances the mapping of chromatin interactions within individual particles. While this enhancement is more significant for cis contacts in closed subcompartments, a subset of ultra-stable trans contacts, including many from active subcompartments, become particularly enhanced by the cryomilling/restriction digestion combination, especially when the cells are not crosslinked before cryomilling.

### Assess the effect of SDS treatment

SDS treatment is commonly used in Hi-C or Hi-C like protocols to improve the accessibility of enzymes to the nucleus and to densely packed chromatin regions, presumably by partially denaturing protein complexes associated with chromatin. However, how exactly the SDS treatment would impact on the mapping of chromatin contacts is not well understood. Unlike in-situ Hi-C, CTCC can gain chromatin accessibility through physical fragmentation of the cell, this affords an opportunity to assess the effect of SDS by performing proximity mapping with and without SDS treatment. CTCC done in the absence of SDS produced a similar contact map to those with SDS (**Supplementary Figure 7a**). Pileup analysis of the local regions of the insulation valleys do not show significant difference in the presence or absence of SDS (**Figure 7a**, left column), indicating that the global structures of TAD and the boundary region between the TADs are generally not affected by the SDS treatment. One the other hand, the loop signal appears to be ‘cleaner’ in CTCC without SDS (**Figure 7a**, right column). Pileup analyses of the DNA breaking distribution relative to the center of the reference nucleosomes show that SDS substantially alters the accessibility of enzymes to these cryomilling generated DNA breakings (**Figure 7b**). First, without SDS, there are two dips on the pileups around 80 bps to the center of the nucleosome (**Figure 7b**, red arrows), which are evident in subcompartments A1, A2, B1 and B2 but not in B3. These dips indicate DNA breaks at these locations are less accessible to mapping enzymes than the surrounding regions without SDS. In the presence of SDS, such dips were absent or smoothed out, indicating that SDS treatment altered the structure of the nucleosomes, presumably by partially dissociating DNA from nucleosomes in these regions. Second, SDS treatment seems to reduce nucleosome position signal across all subcompartments as indicated by the shallower middle valley corresponding to the center position of the nucleosome (**Figure 7b**), suggesting that at least some nucleosome complexes can be disrupted by SDS treatment. By extending the pileup assay of cryomilling generated genomic DNA breakings relative to a number of transcription factors(TF) previously mapped by ChIP-Seq in GM12878 cells (**Supplementary Figure 7b**), it seems that at least for a number of transcription factors, CTCC without SDS can detect some fine structure features at or near the transcription factor binding sites whereas CTCC with SDS tends to smooth out these features, indicating SDS treatment could disrupt or alter the fine, local structure of these regions. CTCC without SDS also shows a higher fraction of short-range contacts and seemingly a steeper pair-frequency-pair-distance curve (**Figure 7c, Supplementary Figure 7c**). These observations suggest that SDS may lead to partial loosening of the chromatin structure thereby disrupting many local and short-range structures, including those mediated by nucleosome and transcription factors, whereas long-range interactions, such as loops seem to be mediated by complexes that are resistant to SDS treatment. Furthermore, CTCC without SDS produces a much lower fraction of trans contacts (**Figure 7d**) which is also observed with micro-C(7, 8). The slightly increased trans signal (same compartment pair) to noise (cross compartment pair) ratio indicates the strategy (pH, ionic strength) we used for dispersing particles without SDS is successful. In all, while SDS is often used to enhance the accessibility to DNA for restriction enzymes, by loosening up the chromatin complex, the same effect can also lead to the disruption of local and fine chromatin structures, and the loose DNA ends resulting from partial denaturation of protein/DNA complexes could also contribute to the noise of proximity mapping.

**Figure 7.**
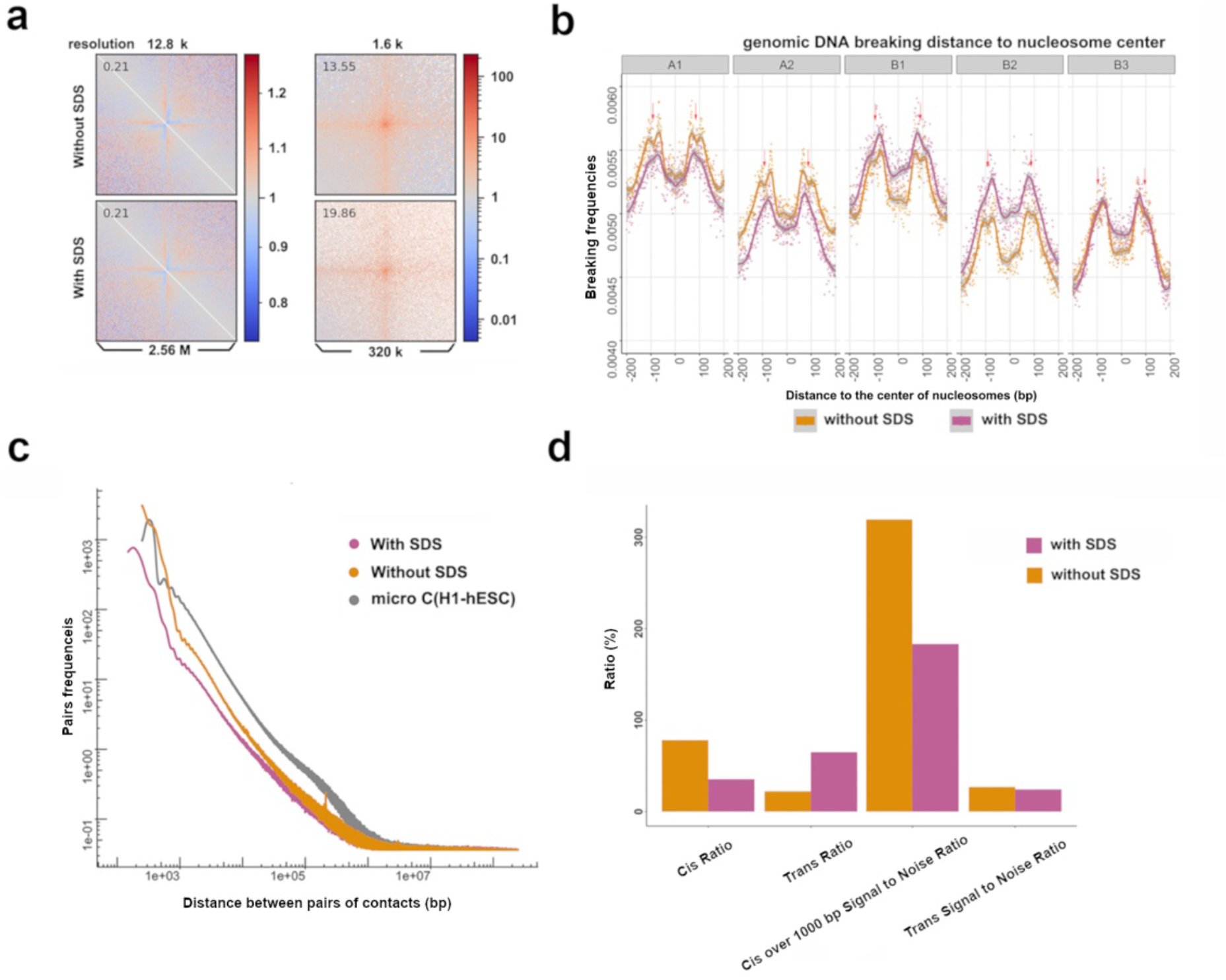
Comparison of CTCC experiments with or without SDS. **a**. Comparison of pileups of contact matrices along the diagonal at local insulation valleys(left) and reference loop positions(right). **b**. Comparison of genomic DNA breakings’ pileup relative to the centers of nucleosomes across different subcompartments of the genome. **c**. Comparison of pair-frequency-pair-distance curve (micro-C data from H1-hESC cells(8) were used as a reference). **d**. Pairs statistics: Cis Ratio = 100 * Cis pair counts / total pair counts, Trans Ratio = 100 * Trans pair counts / total pair counts Cis over 1000 bp Signal to Noise Ratio = 100 * number of “cis same compartment pairs (with Distance more than 1000 bp)” / number of “cis inter compartment pairs (with Distance more than 1000 bp)” Trans Signal to Noise Ratio = 100 * Trans same compartment pairs / Trans inter compartment pairs

### Assess the role of RNA in chromatin structure and interactions

By breaking up the cells into small and open accessible fragments, CTCC provided an opportunity to investigate the role of RNA in chromatin interactions by enzymatic digestion of RNA in the subcellular fragments in-vitro. The effect of RNase A treatment in the CTCC experiment has two aspects. Firstly, RNase A treatment substantially increases the accessibility of enzymes to the heavily transcribed region of the genome. pileup analysis of the position of DNA breaks relative to the center of the nucleosomes’ center shows a clear separation in subcompartment A1 between CTCC experiments with RNase A and CTCC without RNase A, and to a lesser extent in subcompartment A2 (**Figure 8b**), indicating RNase A treatment increases the accessibility of restriction enzymes to actively transcribed genomic regions such as A1 and A2, consistent with several studies showing a predominant association of nascently transcribed RNA to chromatins, especially in highly transcribed regions(37–40). On the other hand, enzyme accessibility to subcompartment B1,B2 and B3 seems to be negatively associated with RNase A treatment (**Figure 8a**), which could be a passive decrease as few RNA are transcribed in these subcompartments, and an increased reads fraction from in the A compartments should accordingly result in decrease in the fraction in the B compartments. Secondly, RNA plays important roles in the structural organization and function of the genome, which is in accordance with previous reports(40–45). RNase A treatment substantially decreased the frequency of contact pairs more than 1kb apart while increasing the ultra-short-range fragments (less than 1kb). (**Figure 8b**), which could be the result from self-ligation, suggesting that the presence of RNA may inhibit self-ligation or that RNA molecules should play a positive role in mediating medium to long range chromatin interactions. Recent research highlighted the importance of transcriptional activity of the A compartment in maintaining the ‘open’ structure of the chromatin(40) and our results also support this point. However, it is interesting to note that even the amount of RNA associated from the A and B compartments is substantially different, RNase A treatment seems to universally decrease cis contacts over 1kb in the B compartments (**Supplementary Figure 8**), indicating a small amounts RNA molecules also play important role in maintaining the structural integrity in these closed subcompartments of the genome.

**Figure 8.**
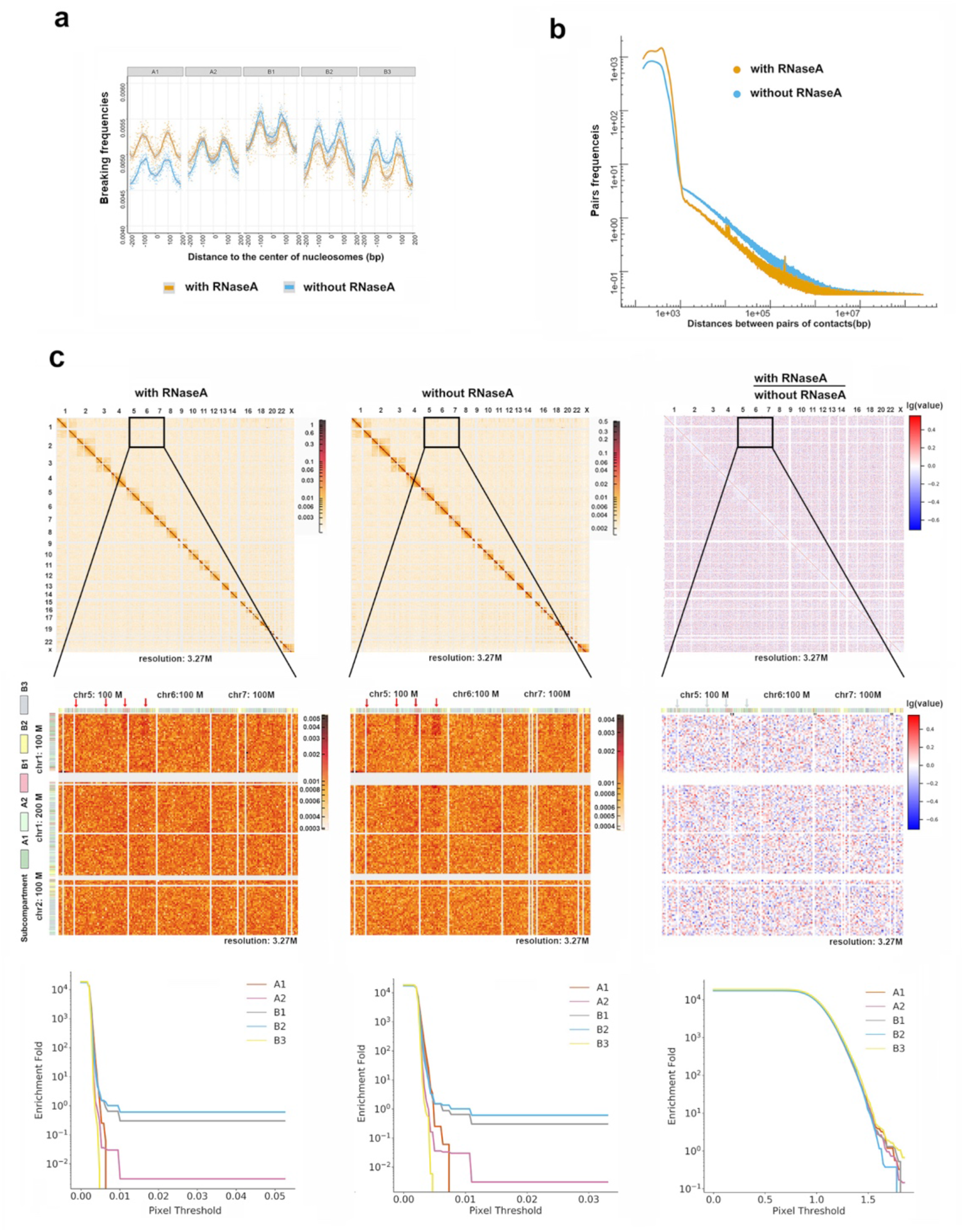
RNase A digestion’s effect on CTCC experiment. **a**. Comparison of cis-pair-frequency-pair-distance curve. **b**. Comparison of genomic DNA breakings’ pileup relative to the centers of nucleosomes across different subcompartments of the genome. **c**. Top panel: Comparison of the contact matrices of CTCC with RNase A (Top panel, left) and CTCC without RNase A (Top panel, middle) and their division (Top panel, right); Middle panel: zoomed in view of a boxed region in the contact matrices; Bottom panel: The trans pixels’ subcompartment enrichment curve at different pixel thresholds for the contact matrics of CTCC with RNase A (left), CTCC without RNase A (middle) and their division matrix (right).

Finally, the ultra-stable trans contacts mentioned above, i.e., those resistant to cryomilling and stable in the in-vitro thawing and crosslinking process, seem not to be affected by RNase A digestion, which is shown not only in the individual contact map (**Figure 8c**, left and middle column, highlighted by the arrow) but also in the subcompartment enrichment curve of the division matrix (**Figure 8c**, right column). This observation, together with the fact that most cis contacts are sensitive to RNase A digestion, suggest that RNA play important roles in maintaining the structural integrity of individual chromosome, whereas the rare but ultra-stable trans contacts between different chromosomes appear be stabilized mostly by protein complexes or factors other than RNA molecules. Thus, although the nascent sites of RNA synthesis are found mostly localized on the surfaces of chromosome territories and between the interface of different chromosomes(39), they don’t seem to contribute to the stable inter-chromosomal contacts observed here in CTCC.

### Relationship between nucleoprotein binding sites and trans contacts

The above analyses show that the certain trans contacts enriched in A1, B1 and B2 subcompartment of the genome observed in CTCC could represent physical contacts that tend to be retained in the same particle after cryomilling, and are stable in-vitro, and seem not be mediated by RNA. This begs the question of who is (are) responsible for such stable inter-chromosome contacts. In the olfactory sensory neurons, Lhx2, EBF and Ldb1 are responsible for bringing OR enhancers from different chromosomes together(46–48), leading to only one OR gene being expressed in that cell. Could it be possible that the stable trans interactions observed with CTCC as discussed above were mediated by or associated with certain nucleoproteins. To address this question, a straightforward yet effective approach is to pile up a bunch of small patches of matrices in the trans region (between chromosomes) of the Hi-C matrix at the ‘crossings’ of the nucleoprotein binding sites as defined by ChIP-seq signals (**Supplementary Figure 9a**). If a particular nucleoprotein is associated with these trans interactions, we should expect to see higher pixel values in the pileup matrix generated according to the binding sites of the nucleoprotein. To do the pileup precisely, we need a high-resolution matrix. However, due to sequencing depth limitations, the enhanced trans patterns seen in the division matrix are only significant at low resolutions and will quickly be buried in noise as resolution increases (data not shown), which is also reflected by the less separation of the subcompartment enrichment curve as the resolution increases (**Supplementary Figure 9b**, right column). This is because the Hi-C matrix is sparse, and the trans region is even more sparse than the cis region, and distribution of the ligation pairs into the local bins will be more dominated randomly as resolution increases, such that the division matrix shows increasing large values with increasing standard deviation (data not shown). We found that the subcompartment enrichment curves for the division matrix of CTCC with HindIII divided by CTCC without HindIII is similar to that of in-situ Hi-C, that high value trans pixels tend to enrich more in A1, B1 and B2 than A2 and B3 subcompartment of the genome (**Supplementary Figure 9b**). It is therefore possible to use previously published, deep sequenced, high-resolution Hi-C data to do the pileup to probe the role of DNA binding proteins in stable trans interactions. Based on the enrichment of stable trans interactions in specific subcompartments as discussed above, we decided to focus on ChIP-Seq peaks coincide with the A1, B1 and B2 subcompartments, and only same subcompartment peaks (i.e., A1-A1, B1-B1 and B2-B2) were paired to generate the coordinates (**Supplementary Figure 9c**). To do this, we wrote two python codes, one for generating the coordinates according to the “same subcompartment ChIP-Seq peaks’ crossings” in the trans region and the other for doing the pileup, including calculating the averaged trans-pileup-matrix (hereafter referred to as trans-pileup-matrix). With these two python codes we pulled the available ChIP-Seq data of the GM12878 cells from the Encode data portal(49, 50) and in-situ Hi-C (MboI) for GM12878 (Lieberman Aiden Lab) mcool file from 4DN data portal(51, 52) to do the pileup. From the all the plotted pileup matrices (**Supplementary Figure 9d**, Supplementary Material), we did not see the classical loop pattern seen in the diagonal of the Hi-C matrix, which were known to be mediated by CTCF and cohesin(6,53–56) and the fine loop pattern seen in ultra-resolution micro-C matrix, which was prove to be mediated by transcriptional activity(7). We then calculated the pileup-matrix-pixel-value-sum (hereafter referred to as pileup-sum) which is the sum of the pixel values for the trans-pileup-matrix. We found the pileup-sum values for different nucleoproteins from the A1-A1 region generally have higher pixel values than those from B1-B1 or B2-B2 regions (**Supplementary Figure 9e**), which is in accordance with our previous observations. The pileup-sum values are not correlated with the total coordinate number for the individual nucleoproteins (**Supplementary Figure 9f**), indicating the differences in pileup-sum could be caused by where the individual nucleoproteins bind on the reference genome. Using the UMAP algorithm(57), pileup matrices from different nucleoproteins’ ChIP-Seq peaks could be categorized into four clusters (**Supplementary Figure 9g**), which are related to the subcompartment binding properties of the nucleoproteins (such as those only bind to A1 subcompartment of the genome, i.e., strict A1 binder) and pileup-sum. It is interesting to note that nucleoproteins who have higher pileup-sum in A1 subcompartment generally have higher pileup-sum in B1 subcompartment, which is shown by the correlation matrix (with a pearson correlation score of 0.46) (**Supplementary Figure 9h**), indicating these nucleoproteins should have a general ranking of the ability of bringing chromosomes together, irrespective of chromatins’ depression (activation) status. According to the value of pileup-sum of each nucleoprotein for each subcompartment of A1, B1 and B2, we could rank the nucleoproteins (**Supplementary Figure 9i**). As a reference, CTCF, which is a known transcription factor participating in the TAD formation(4,6,54,55), only ranked medium in both A1 and B1 pileup-sum (**Supplementary Figure 9i**, blue arrow), indicating a lower likelihood of being related in these trans interactions seen in Hi-C and CTCC. On the other hand, FOS, a protein important in lymphocyte development(58, 59), rank top in all three subcompartment. Interestingly, JUNB and JUND, two known FOS partners, ranks low in all three subcompartments (**Supplementary Figure 9i,** black dashed line boxes), indicating other partners may be associated with FOS to mediate the trans interactions.

## Discussion

In this study, we developed CTCC, a method combining cryomilling and TCC, for the mapping of chromatin interaction in cells or tissues. At the global level, cryomilling shows unbiased physical breaking of DNA across the entire genome (**Figure 3a**), but at the local level it appears sensitive to fine structures formed by strong protein-DNA interaction such as the nucleosome complexes, suggesting a potential to combine cryomilling with high resolution mapping techniques such as MNase to probe fine protein/DNA complexes structure in the nucleus. The unbiased physical breaking of the nucleus results in enhanced chromatin access, especially when combined with enzymatic digestion of DNA in the cryomilled cell particles, and this access enhancement leads to more even and efficient sampling of chromatin interactions across the entire genome, especially in the closed and more compact subcompartments.

Because CTCC maps chromatin interaction in a large population of subcellular fragments, a key question that needs to be addressed before establishing CTCC as a valid chromatin conformation analysis technique is if CTCC would be able to capture global chromatin conformation features of the entire cells. A second and perhaps equally important question is if some chromatin interactions are disrupted by the cryomilling process and if different subtypes of chromatin interactions are differently affected by this disruption. In this study, we carried out CTCC of GM12878, a model cell line widely used in the method development of Hi-C and related genomics methods(2,3,10,60,61). Our results reveal a DNA contact map that is similar to those obtained by Hi-C, suggesting that detection of DNA proximity in cellular fragments randomly generated by cryomilling can reconstitute the ensemble global chromatin conformation features. This is a significant finding because it suggests that chromatin conformation analysis could be applied to tissues and solid organs using the relatively simple and more accessible CTCC protocol as compared with GAM.

To address the second question, we carried out CTCC mapping of chromatin interactions in GM12878 cells under different conditions and compared the results between CTCC and Hi-C and between different CTCC experiments. In principle, Hi-C, in which proximity ligation is carried out in-situ inside the nucleus, should detect both physical contact and proximity in space without physical contact (**Supplementary Figure 4a**), since in both cases, proximity information would be captured. In CTCC, cryomilling should preferentially dissociate loci that are not physically associated (**Supplementary Figure 4a, A** and **B**) with each other and should therefore enrich physical contact (**Supplementary Figure 4a, C** and **D**). Meanwhile, the enormous number of subcellular particles created by extensive cryomilling also create strong competitive inter-particle ligation events, which could increase noise. The result of CTCC should be the competition between intra and inter particle ligation events. Compared with conventional and in-situ Hi-C, CTCC enhances the mapping of cis contacts in the closed compartment of the genome, which should be due to the enhanced chromatin access mentioned above. The trans contact signal, however, is substantially reduced in CTCC, which also shows higher background noises as indicated by higher cross-subcompartment ligations in CTCC. Further analyses reveal that active subcompartments A (A1 and A2) are more prone to such cross-subcompartment noises than the inactive B subcompartments (B1, B2 and B3 cis contacts). A major source of cross-subcompartment noises may be DNA ligation between different subcellular particles. Active subcompartments of the genome represent extended, unfolded chromatin fibers that take more space than other subcompartments of the genome in the nucleus. Consequently, cryomilling could generate more subcellular particles from the active subcompartment than the inactive subcompartments, leading to more inter-particle ligation events and substantially increased cross-subcompartment ligation noise (**Supplementary Figure 4b**). The comparative analyses between Hi-C and CTCC also reveal that while most true trans signal (i.e., same subcompartment trans contact) are weak and unstable in CTCC (such as those diminished same subcompartment trans contacts in the B3 subcompartment), some trans contacts (such as those cryomilling resistant trans contacts in B1) are stable and can be efficiently mapped in CTCC.

The stabilities of chromatin interactions can also be analyzed by CTCC with different crosslinking protocols, namely crosslinking of intact cells before cryomilling (in-situ crosslinking CTCC, crosslinking before cryomilling) or crosslinking of fragmentized cell particles after cryomilling (in-vitro crosslinking CTCC, crosslinking after cryomilling). While large scale chromatin structures such as TAD can be efficiently captured by in-vitro crosslinking CTCC, a large amount of short to medium range chromatin contacts, including the fine structural features of inter-nucleosome interactions, require formaldehyde crosslinking of the cells before cryomilling. Moreover, the in-vitro crosslinking CTCC captured much less inter-chromosomal (trans) contacts as compared with the in-situ crosslinking CTCC, suggest that subcellular regions made of DNA from different chromosomes are unstable in the cryomilling process or during crosslinking in-vitro with formaldehyde in solution. On the other hand, while most trans contacts are unstable hence unobservable in the in-vitro crosslinking CTCC, trans contacts between certain loci of the genome are surprisingly stable and become much more standing out from the overall reduced trans signals in the in-vitro crosslinking CTCC. While these differences are similar to those seen between Hi-C and CTCC, the comparison between the two CTCC experiments directly reveal the effect of formaldehyde crosslinking in the stabilization and capture of a significant amount of chromatin contacts. This observation justifies the use of formaldehyde crosslinking in chromatin conformation capture studies(16, 17). However, as discussed earlier, a potential problem of this in-situ crosslinking approach is the slow diffusion of formaldehyde across the entire cell volume, resulting in different crosslinking efficiency in different subcellular regions. With CTCC, the chromatin conformation is captured by flash freezing. Physical breaking of the cells under cryogenic conditions ensures the preservation of native chromatin interactions in cryomilled subcellular fragments. Although in the current CTCC protocol the cryomilled cell fragments are also stabilized by chemical crosslinking for subsequent DNA proximity mapping, the crosslinking of subcellular fragments could be more rapid, efficient and less diffusion biased. Moreover, the crosslinking reagents could be extended from formaldehyde to other chemicals that may not be amenable to in-situ studies but more efficient for in-vitro capturing protein complexes mediating the chromatin interactions(62, 63).

Chromatin interaction mapping in cryomilled particles can be enhanced by restriction digestion of the DNA in individual particles, presumably by generating additional DNA ends that can facilitate proximity ligation. Since the increased DNA digestion occur mostly within individual particles due to the penetration of restriction enzymes into the cryomilled cell particles rather than between different particles, most of the enhanced proximity ligation events occur between loci of the same genomic subcompartments, i.e., HindIII digestion on cryomilled cell fragments enhance preferably the detection of true chromatin contacts but not the noise. Since most of the cryomilled subcellular particles contain DNA from the same chromosome, the most significant mapping enhancement (increase of mapped contact frequency) by restriction digestion of the cryomilled particles is seen with cis contacts. Some subcompartments (e.g., A2 and B3) show more significant mapping enhancement than others (e.g., A1), probably because DNA access by restriction enzymes in different subcompartment is differently enhanced by cryomilling. As discussed above, a small fraction of cryomilled subcellular particles composed of inter-chromosomal contacts are stable and resistant to cryomilling. The mapping of these ultra-stable trans contacts are also greatly enhanced by restriction digestion of the cryomilled subcellular particles, in a way similar to the mapping enhancement seen with the cis-contacts. Interestingly, pileup analyses for local features around insulation score valleys and loops do not show significant difference with or without restriction digestion of the cryomilled subcellular particles. This observation raises an intriguing question if these structural features could be captured by ligation of DNA breaks directly generated by cryomilling. In other words, DNA ends corresponding to the insulation boundary and loops occur mostly on the surface of cryomilled cell particles. Further studies will be required to address this question.

Further analyses of trans regions in the contact frequency map show that many of the trans contacts enhanced by cryomilling/HindIII digestion belong to A1, B1 and B2 subcompartments while most of the trans contacts of the A2 and B3 subcompartment are unstable. As discussed above, different crosslinking protocols (i.e., in-situ and in-vitro crosslinking) provide a means to differentiate subcellular particles that are stable or not in cryomilling or subsequent in-vitro thawing and crosslinking, HindIII digestion further enhances the mapping of chromatin interactions in those stable particles. As a result, division of the matrix of “crosslinking after cryomilling with HindIII digestion” by that of “crosslinking before cryomilling without HindIII digestion” shows that ‘crosslinking after cryomilling’ and ‘HindIII digestion’ synergistically enhance those ultra-stable trans signal that belong to the A1, B1 and B2 subcompartments. Thus, if the mapping focus is on the stable inter-chromosomal interactions, the specific CTCC protocol should be crosslinking after cryomilling followed by HindIII digestion of the cryomilled cell particles. A plausible explanation for the above observations could be that certain loci belonging to A1, B1 and B2 subcompartments form entangled trans contacts that are so stable that they tend to remain with each other in individual particles during the cryomilling and subsequent in-vitro crosslinking processes, while most inter-chromosomal contacts belong to the A2 and B3 subcompartments tend to fall apart, either in the cryomilling or during the in-vitro crosslinking process. If cells were crosslinked before cryomilling, either entangled (CTCC stable) or loosely (CTCC unstable) associated chromosome territories would be crosslinked inside the cell and remain together either in the cryomilling and/or subsequent in-vitro proximity ligation processes (**Supplementary Figure 6d**). As a result, CTCC with in-situ crosslinking and conventional Hi-C would not be able to effectively distinguish these two structures. This model also can explain why CTCC with in-situ crosslinking produced much higher overall trans signal and relatively lower cis signal than in-vitro CTCC, and that division of the corresponding matrix of ‘crosslinking after cryomilling’ by those of ‘crosslinking before cryomilling’ produced overall higher cis (> 1) and low trans values (< 1) (**Supplementary Figure 6c upper panel**). CTCC with and without SDS provided further insights into the stability of the chromatin interactions and potential impact of the use of harsh detergent in chromatin interaction mapping. These comparative analyses indicate that most of the global structural features such as TADs and boundary regions are generally not affected by the SDS treatment, although the loop signal appears to be cleaner in CTCC without SDS. However, SDS treatment apparently can cause partial loosening of the chromatin structure thereby disrupting many local and short-range structures, including those mediated by nucleosome and transcription factors, whereas long-range interactions, such as loops, seem to be mediated by complexes that are resistant to SDS treatment. Our analyses also raise an important question that the large fraction of trans contacts commonly observed in conventional Hi-C with SDS treatment may be caused, at least partly, by partial denaturing of chromatin complexes and releasing DNA strands in the crowded nuclear environment thereby increasing trans ligation.

Finally, CTCC with and without RNase treatment provided insights into the role of RNA in chromatin organization. In the active subcompartment A1 and A2, RNase treatment enhanced the DNA accessibility, probably by removing the excessive amount of nascent RNA transcripts that may hinder the restriction digestion and proximity ligation of DNA. Meanwhile, RNase A treatment substantially decreased the frequency of contact pairs more than 1kb apart while increasing the ultra-short-range fragments (less than 1kb, or self-ligation). It is well established that non-coding RNA plays important roles in the regulation of gene expression, for example, some long non-coding RNA molecules have been shown to play an active role in the repression of genes in certain chromatin regions of the repressing structures as part of their functional mechanisms.(5,42,64). Also, recent research shows that active transcription is key in maintaining the open structure of the A compartment(40). Our results are thus consistent with these observations. However, RNaseA’s negative effect on the cis medium to long range contacts are not limited in the two open subcompartments(A1 and A2) but also in the other three closed subcompartments(B1, B2, and B3), indicating a general role RNA play in maintaining the structural integrity of the genome.

Surprisingly, the ultra-stable trans contacts that are not only resistant to cryomilling and stable in the in-vitro thawing and crosslinking process but are also not affected by RNase A digestion. Our comparative analyses between CTCC with and without RNase treatment suggest that the ultra-stable trans contacts detected by CTCC are not sensitive to RNase A treatment, This is an interesting finding given the fact that the nascent sites of RNA synthesis are found mostly on the surfaces of chromosome territories and between the interface of different chromosomes(39). Apparently, these nascent RNA molecules are not involved in stable inter-chromosomal interactions.

To further address the question of who is responsible for these CTCC stable, RNase A resistant inter-chromosome contact, we conducted extensive trans pileup according to known binding position of various nucleoproteins. These analyses suggest that certain DNA binding proteins have a higher tendency to be involved in the stable trans inter-chromosomal interactions. Interestingly, there seems to be a correlation between the ability of nucleoproteins mediating trans A1-A1, and B1-B1 contacts, i.e., those having higher potential mediating A1-A1 trans contact generally also have higher potential mediating B1-B1 trans contact, and vice versa. Interestingly, many transcription factors such as the GATA family (GATA1-6) are known to both active and repress gene expression through long range chromatin interactions, probably by mediated stable trans interactions in both the active and repressive compartments as discussed above(65).

In summary, our studies establish CTCC as a novel and valid chromatin mapping technique that complements many existing methods such as Hi-C, GAM, SPRITE etc. Since cryomilling can be easily adapted to solid organs and tissues, CTCC presents a readily accessible and efficient method for chromatin conformation analyses in solid organs and tissues. By performing CTCC under different conditions, our studies enable detailed analysis of the stability of different subtypes of chromatin interactions and associated protocol parameters. These systemic and comparative analyses not only help to better understand the chromatin conformation mapping approach itself but also provide new insights into the stability and assembly mechanisms of chromatin organization.

---

**Supplementary Figure 1.**
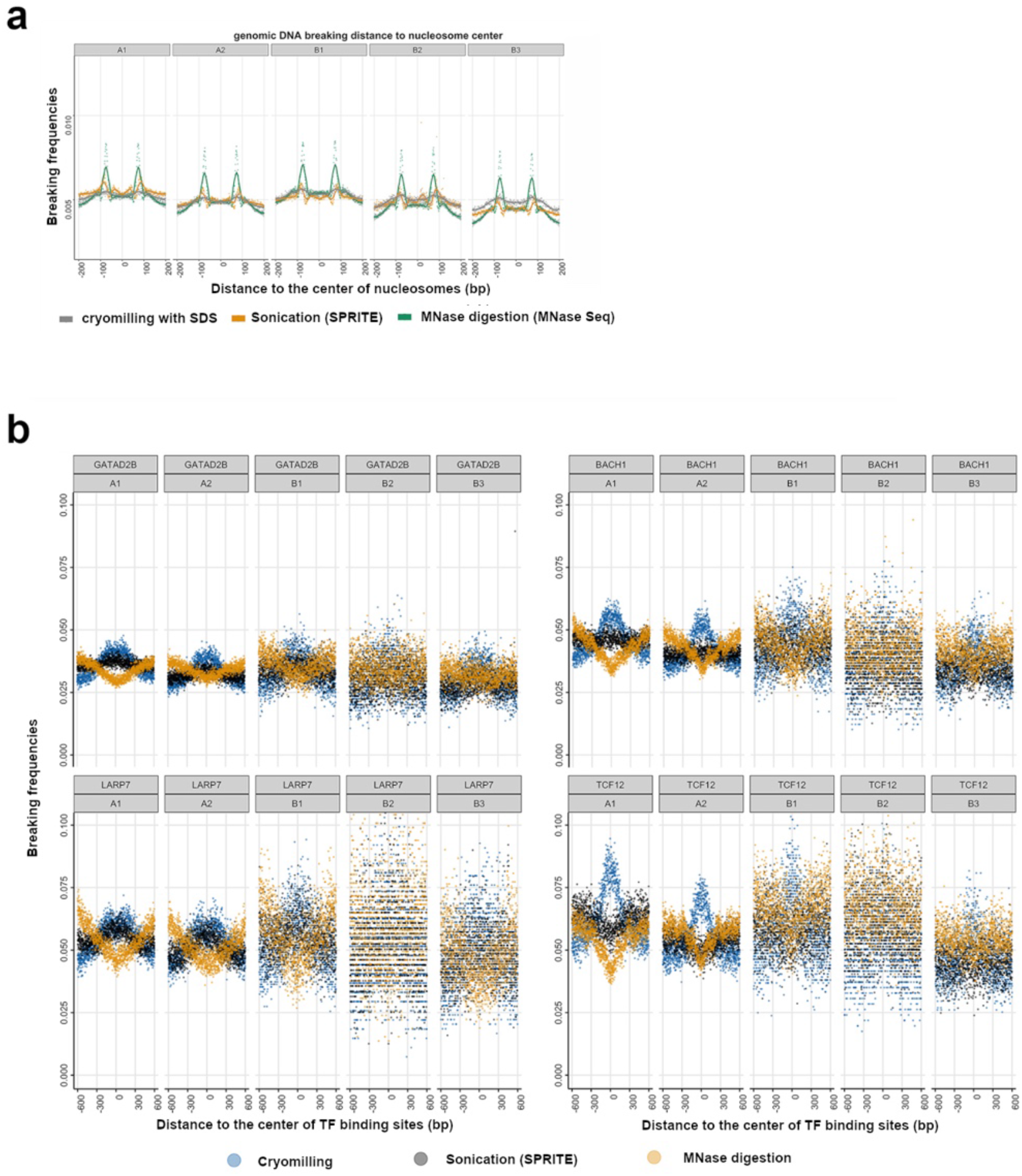
Comparison of cryomilling with sonication, and MNase digestion in terms of DNA breaking positions to various nucleoprotein binding sites. **a.** Comparison of pileups of DNA breaking positions relative to the center of nucleosomes. **b**. Comparison of pileups of DNA breaking positions relative to the binding position of transcription factors (4 typical patterns are shown here).

**Supplementary Figure 2.**
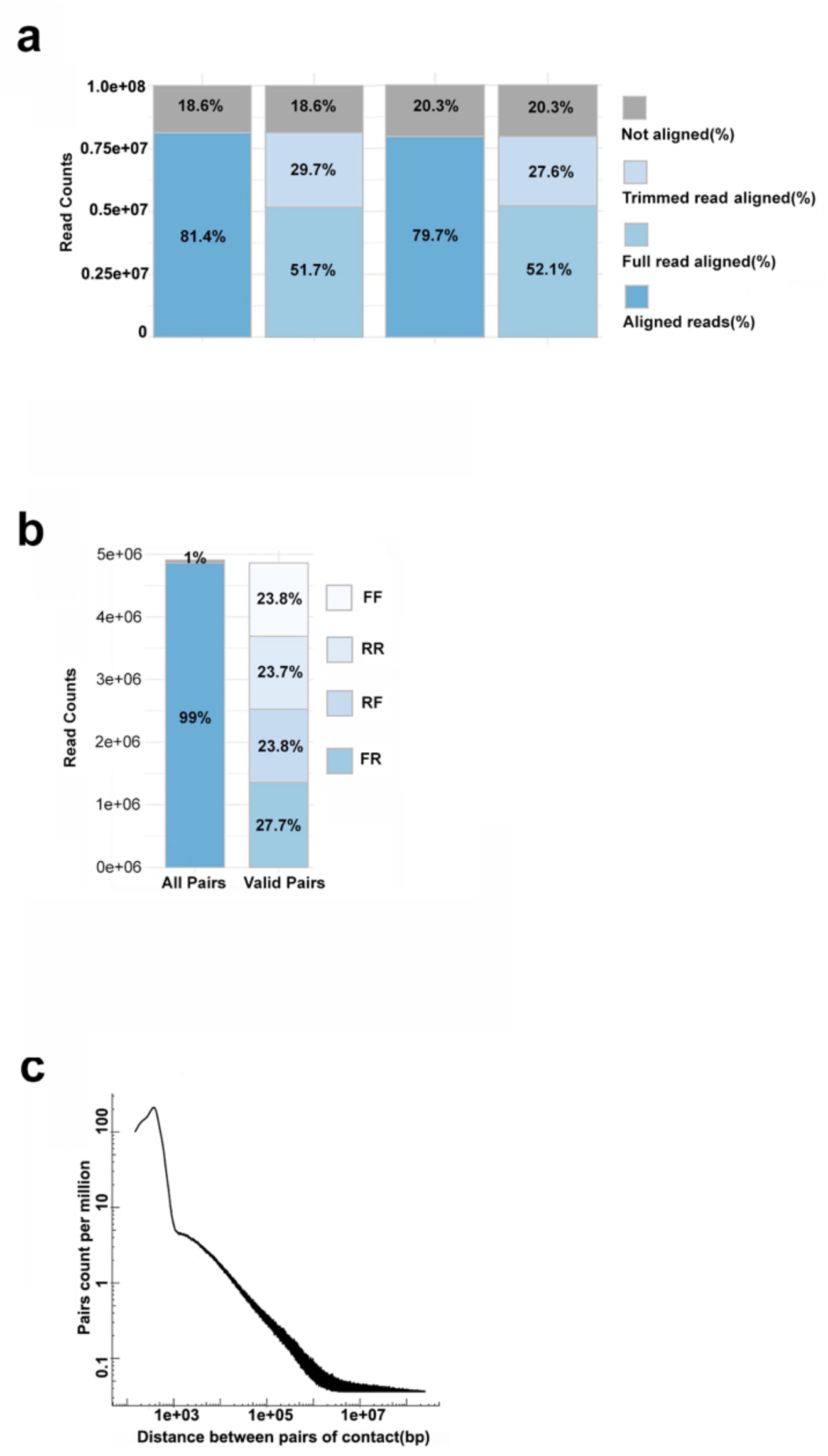
Mapping statistics of a typical CTCC experiment, i.e., CTCC with crosslinking in-vitro (crosslinking after cryomilling) with HindIII digestion. a: Mapping statistics). b: Pair statistics. c: Pair-frequency-pair-distance curve.

**Supplementary Figure 3 a.**
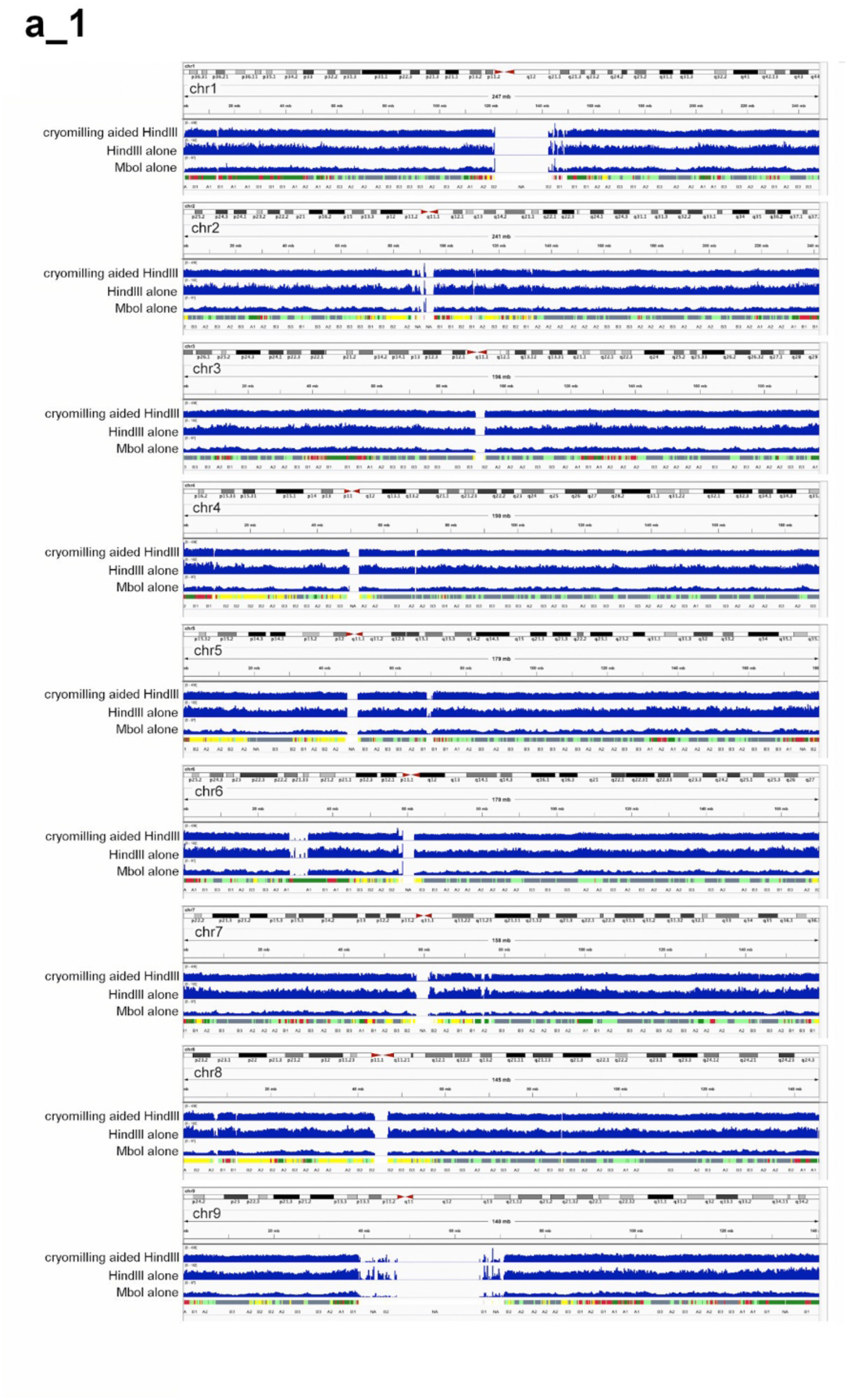
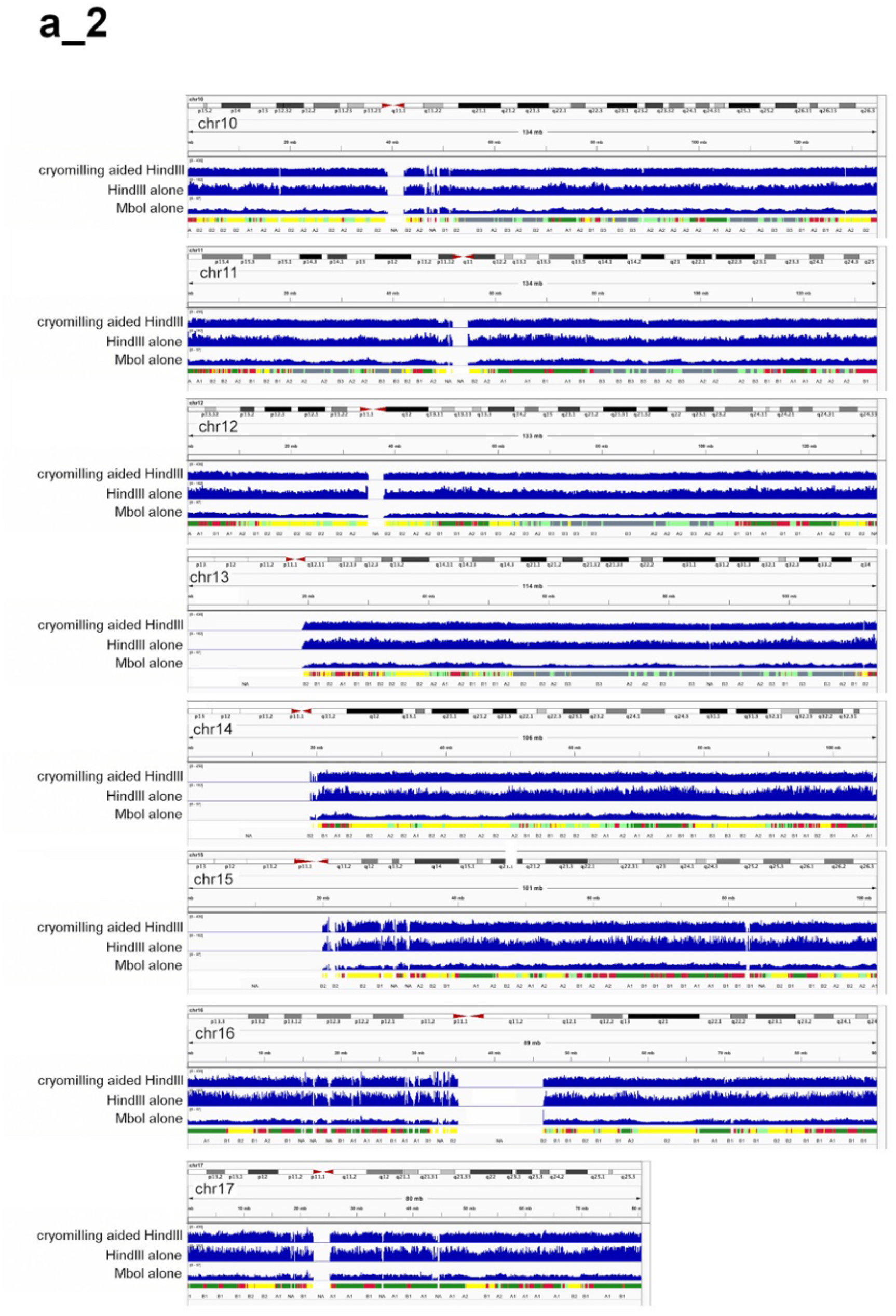
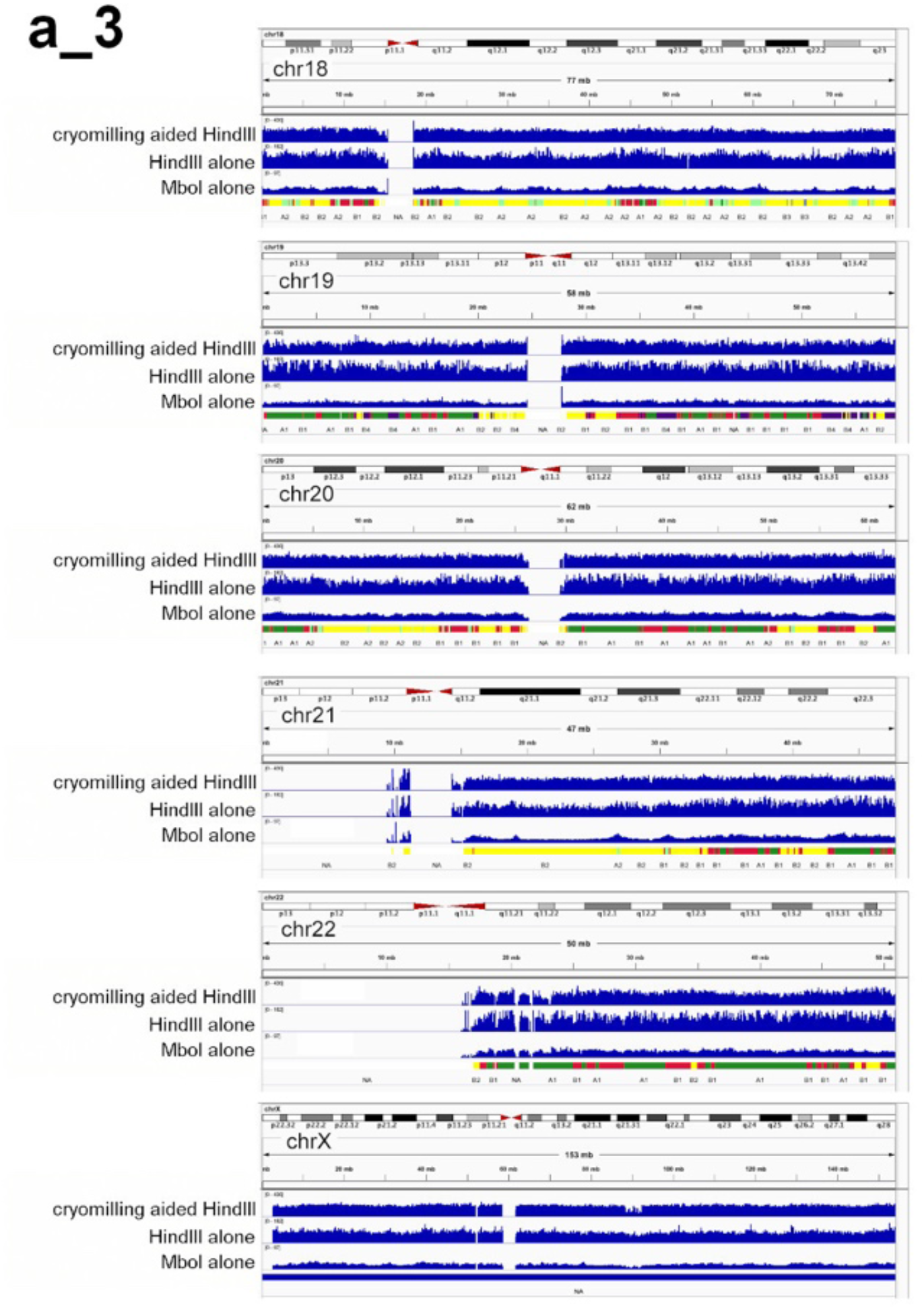
Genome wide comparison of cryomilling aided HindIII digestion and HindIII, MboI digestion alone.

**Supplementary Figure 3 b.**
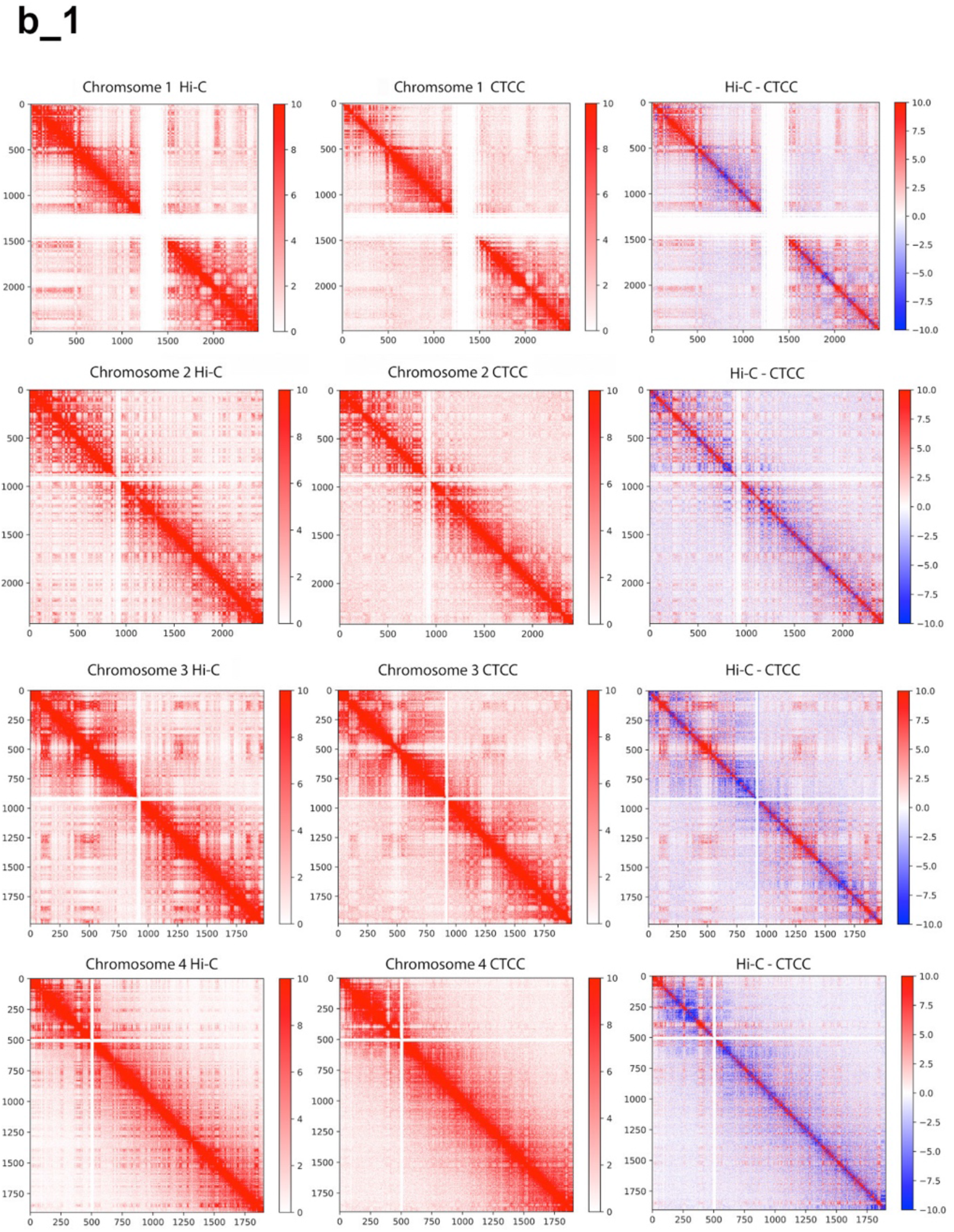
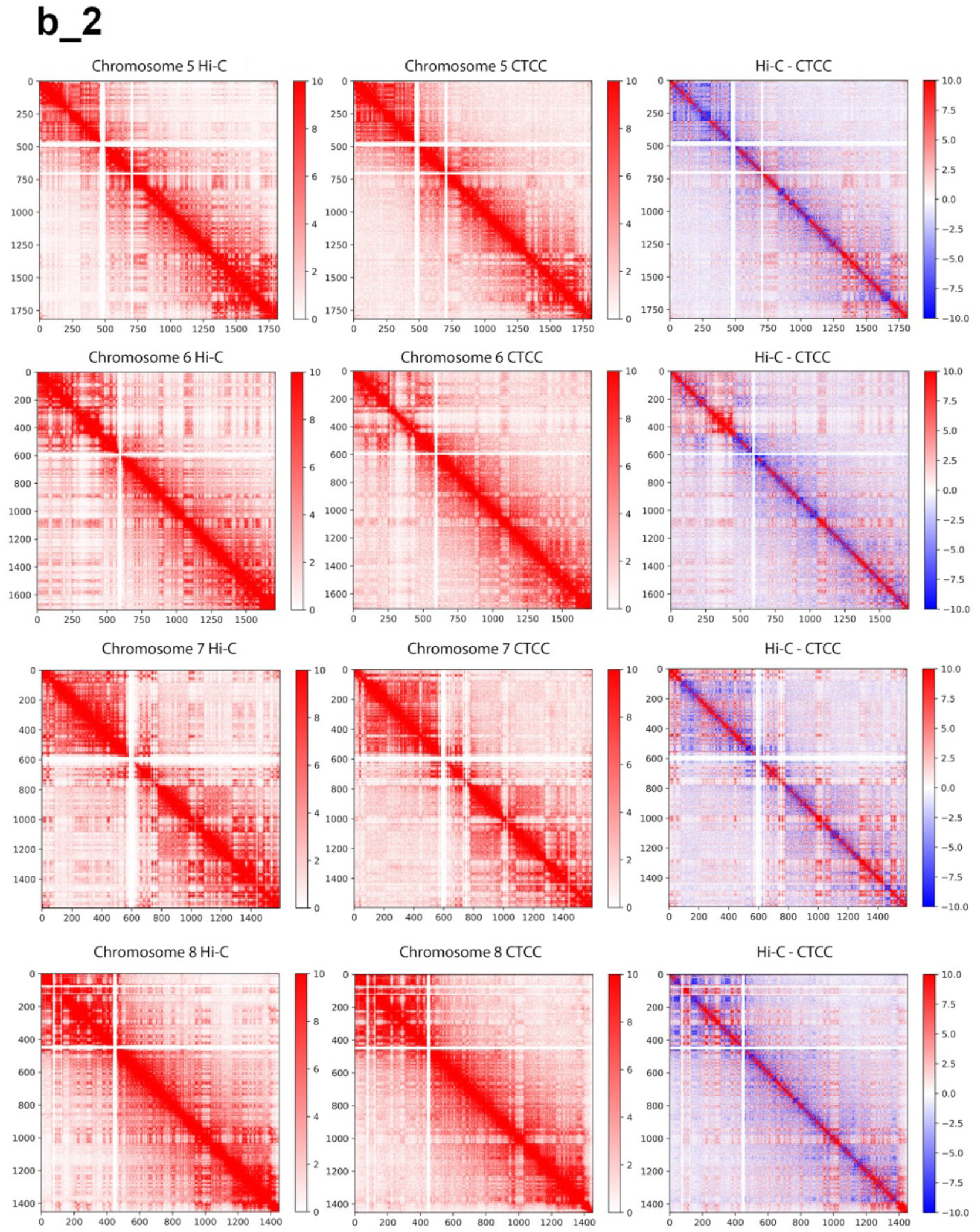
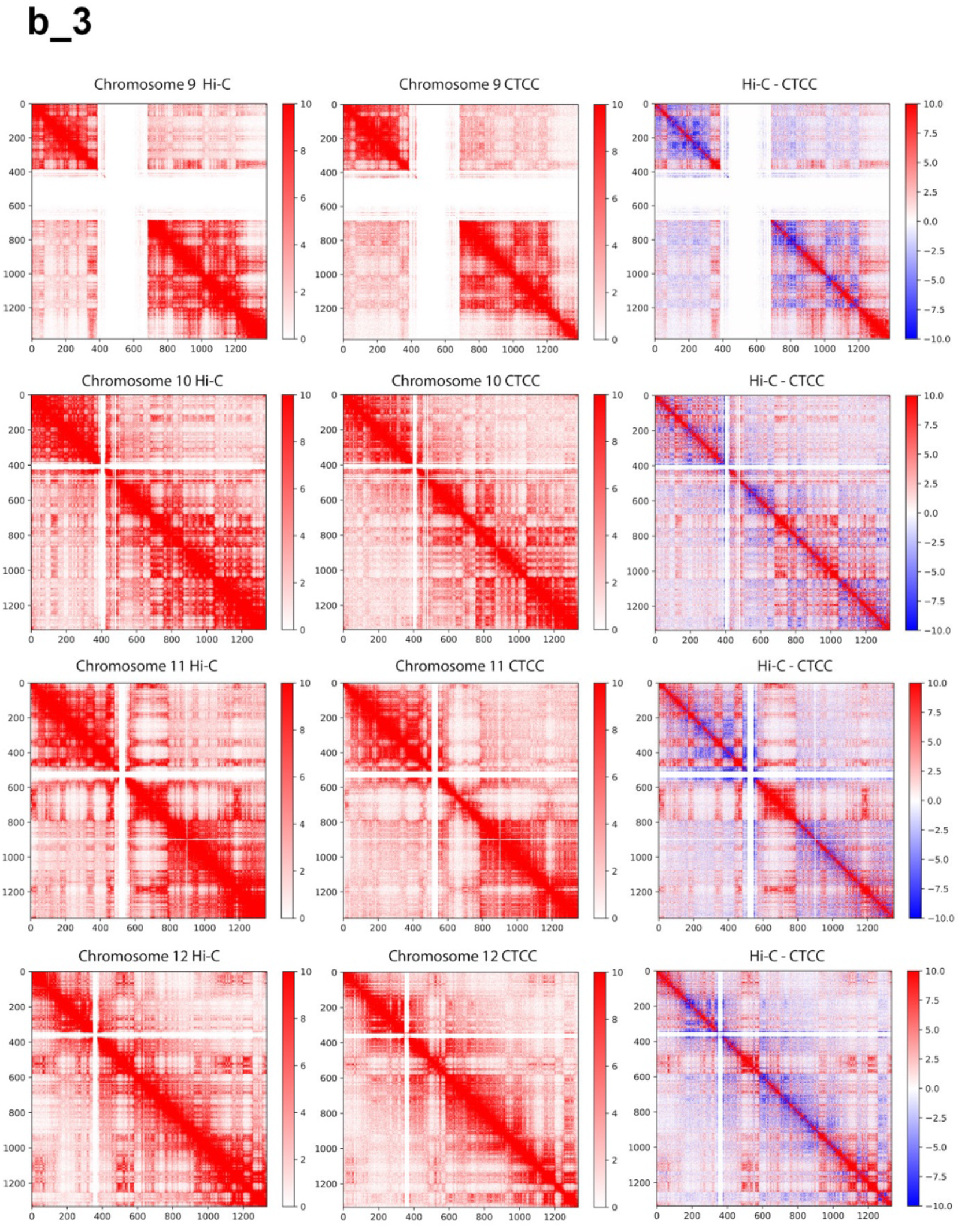
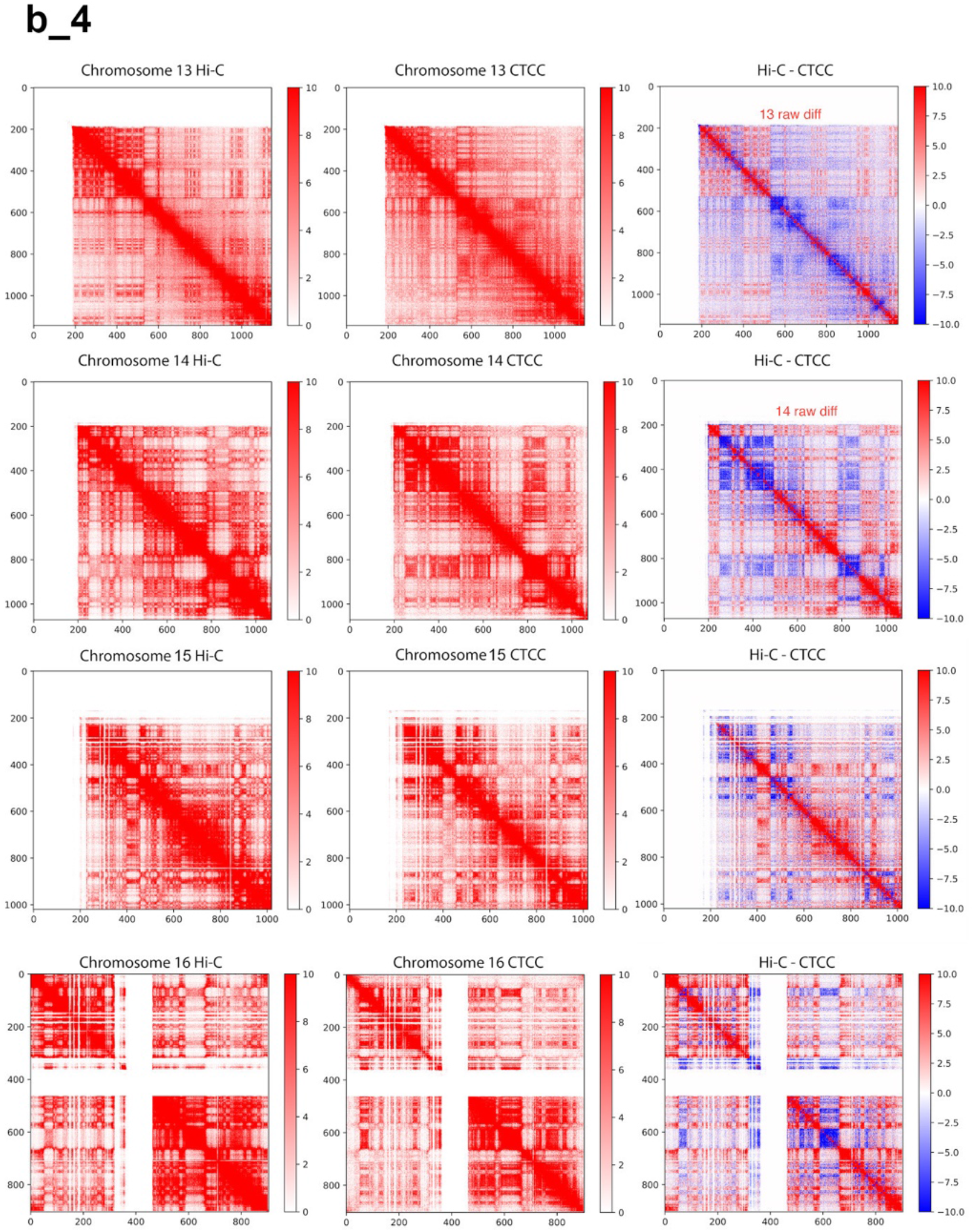
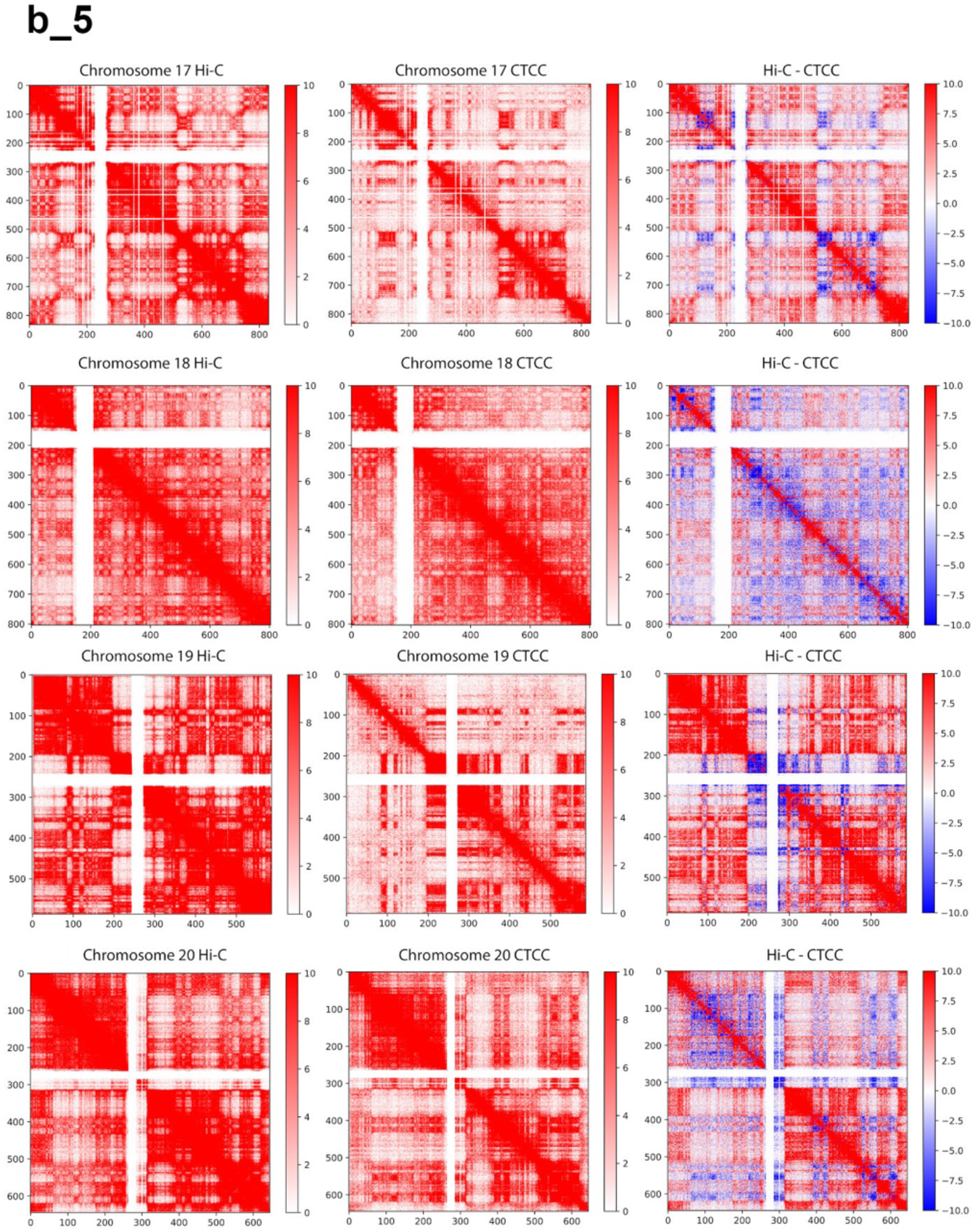
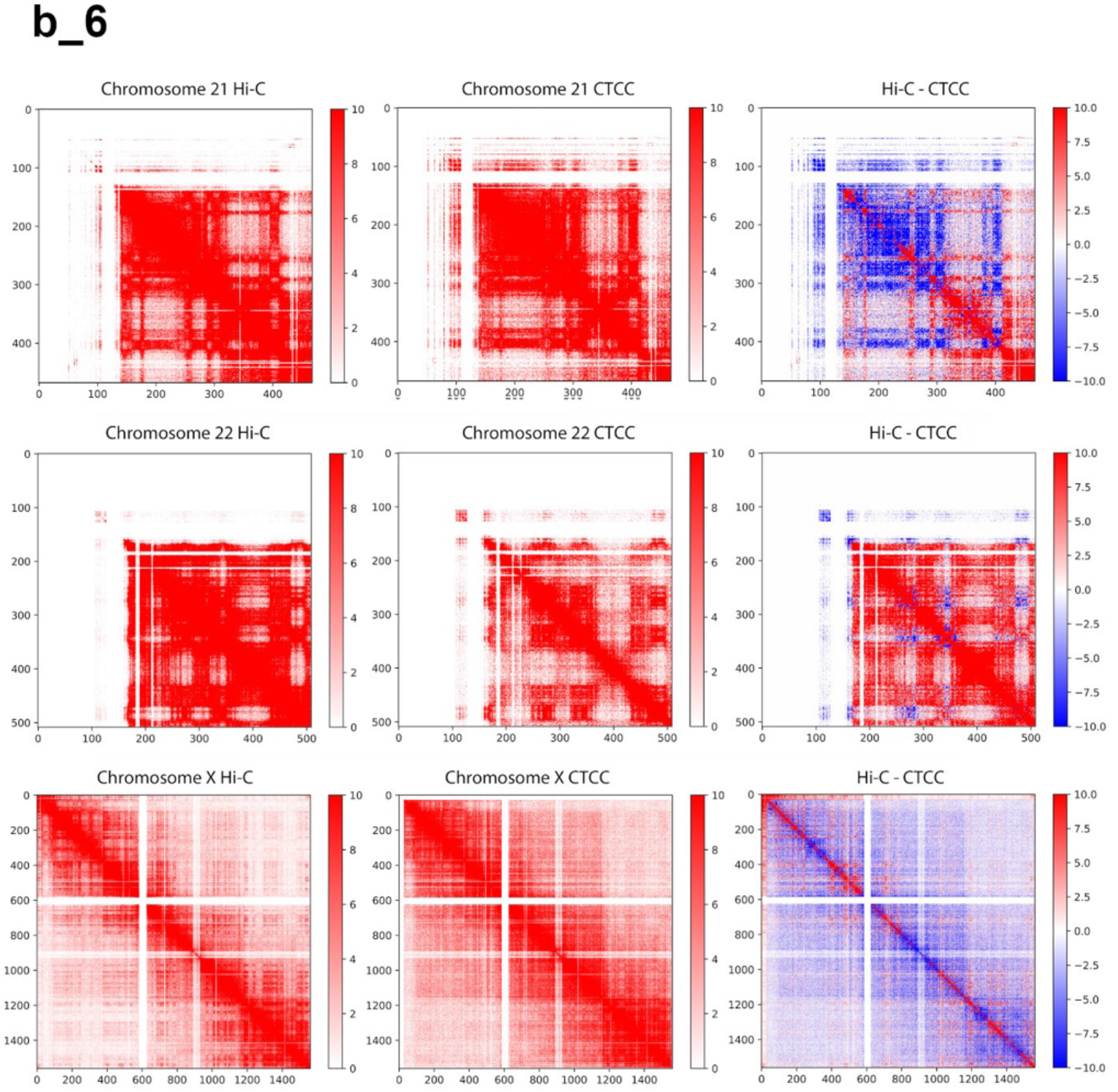
Contact matrices of CTCC with HindIII, dilution Hi-C with HindIII and subtraction of the two across different chromosomes.

**Supplementary Figure 3 c.**
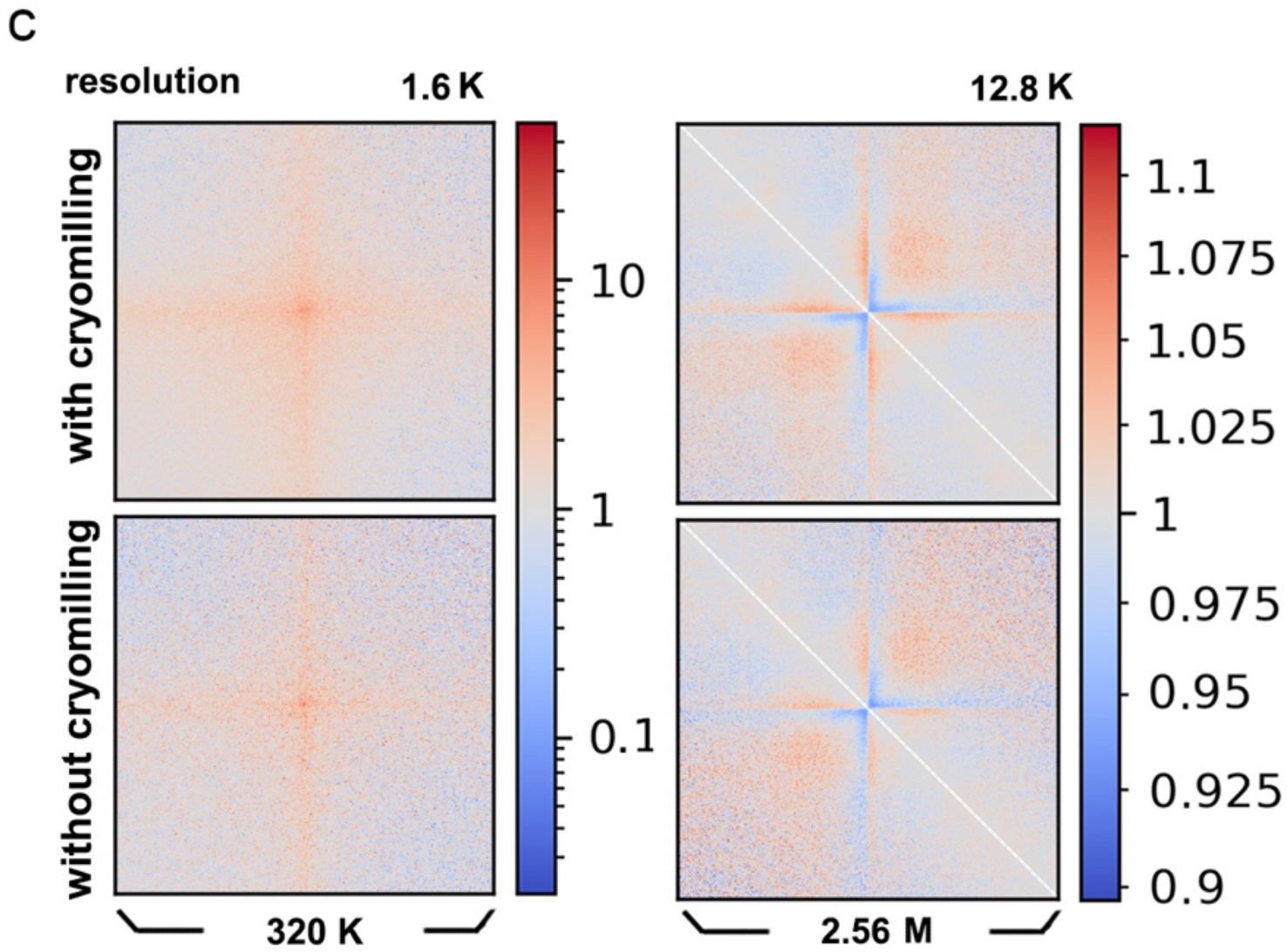
Comparison of pileups of matrices’ local features along the diagonal of the contact matrix at insulation score valleys (between neighboring TAD domains) and at known loop positions for in-situ Hi-C (without cryomilling) and CTCC with HindIII (with cryomilling)

**Supplementary Figure 4.**
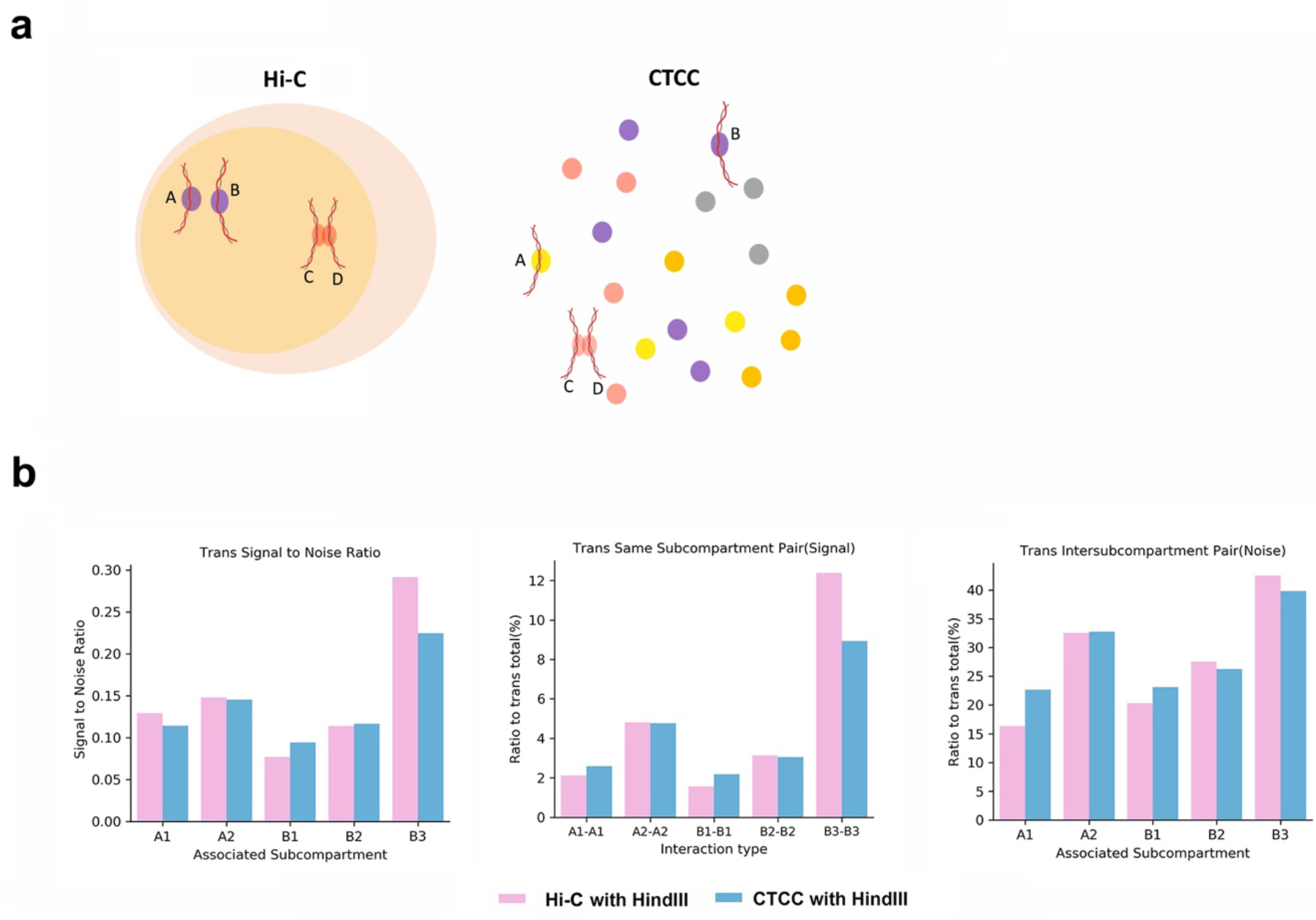
**a.** Principal illustration of two kinds of “contacts”, physical contacts and proximity in space that could both captured in Hi-C but could be differentiated in CTCC **b.** Comparison of raw trans pairs statistics between dilution Hi-C (diHi-C with HindIII) and CTCC (CTCC with HindIII) in terms of “Signal to Noise Ratio”, “Signal Ratio” and “Noise Ratio”

**Supplementary Figure 5.**
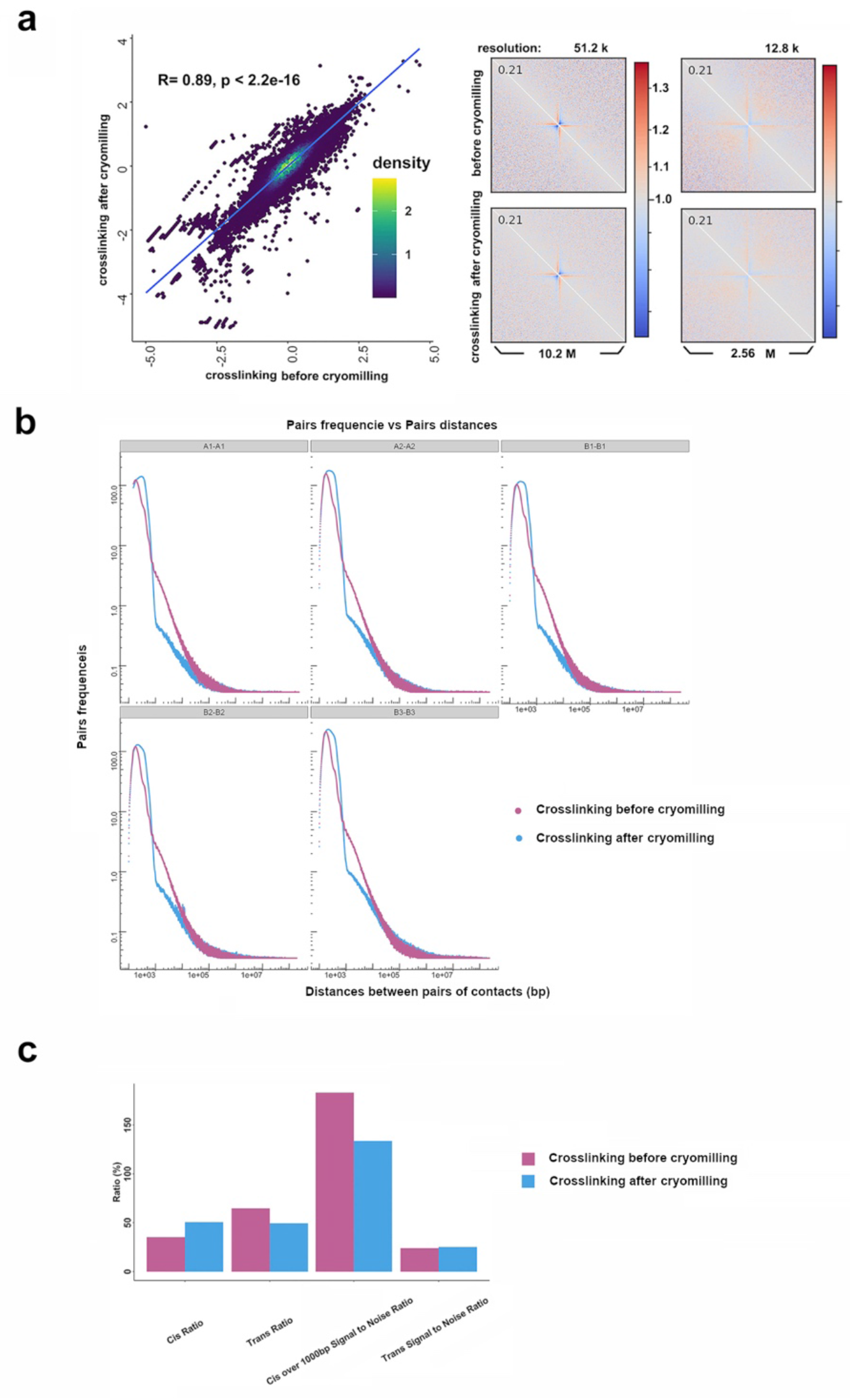
Comparison of the effect on CTCC experiment of two crosslinking strategies, crosslinking before cryomilling and crosslinking after cryomilling. **a**. Left panel: Pierson correlation of insulation score at intermittent locations across the genome between two crosslinking strategies. Right panel: Pileup of local insulation score valleys (the bordering region between TAD domains) along the diagonal of the contact matrix. **b**. Same compartment pair frequency-pair distance curve. **c**. Raw pair statistics including cis pair ratio, trans pair ratio, cis over 1000 bp signal to noise ratio, trans signal to noise ratio.

**Supplementary Figure 6.**
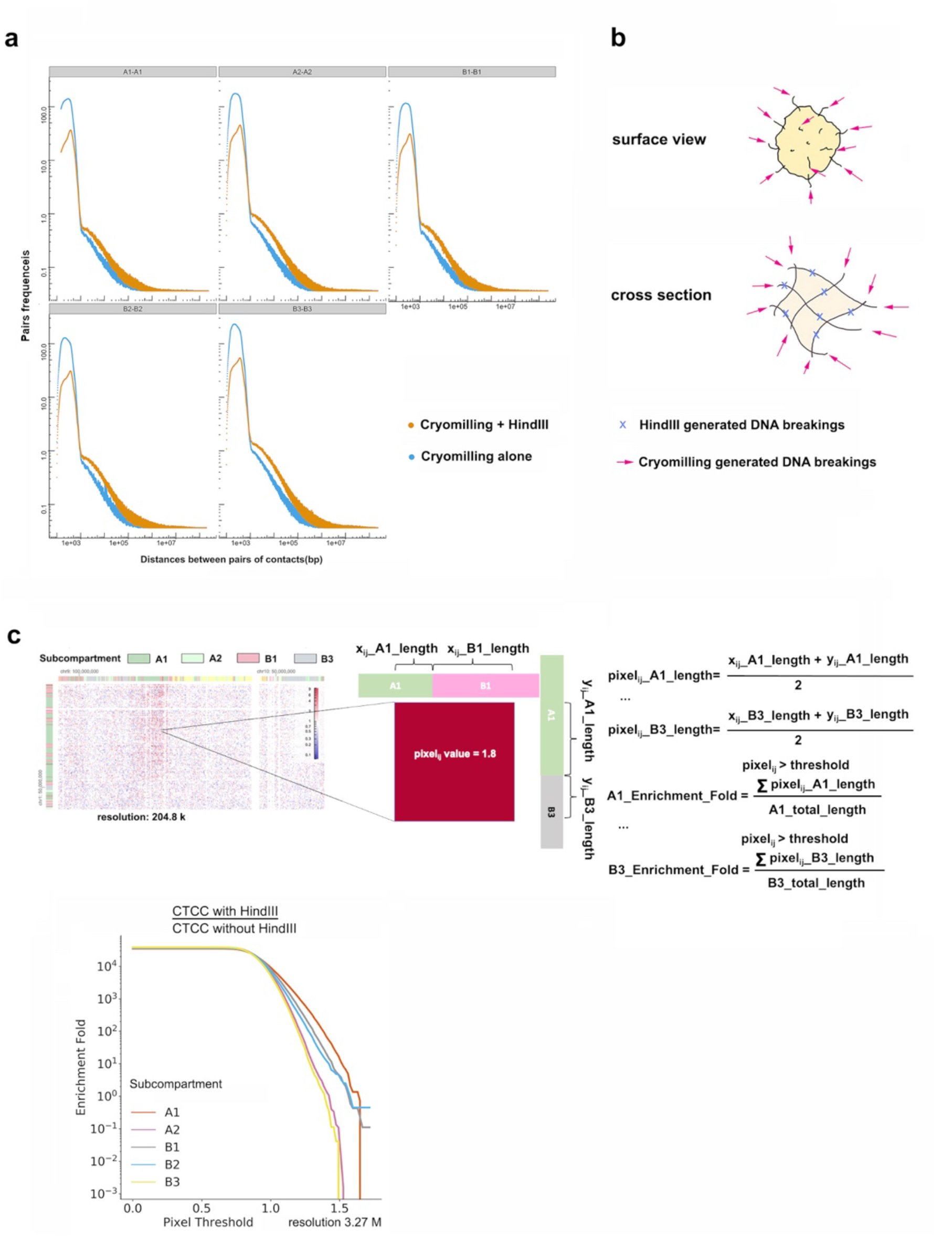
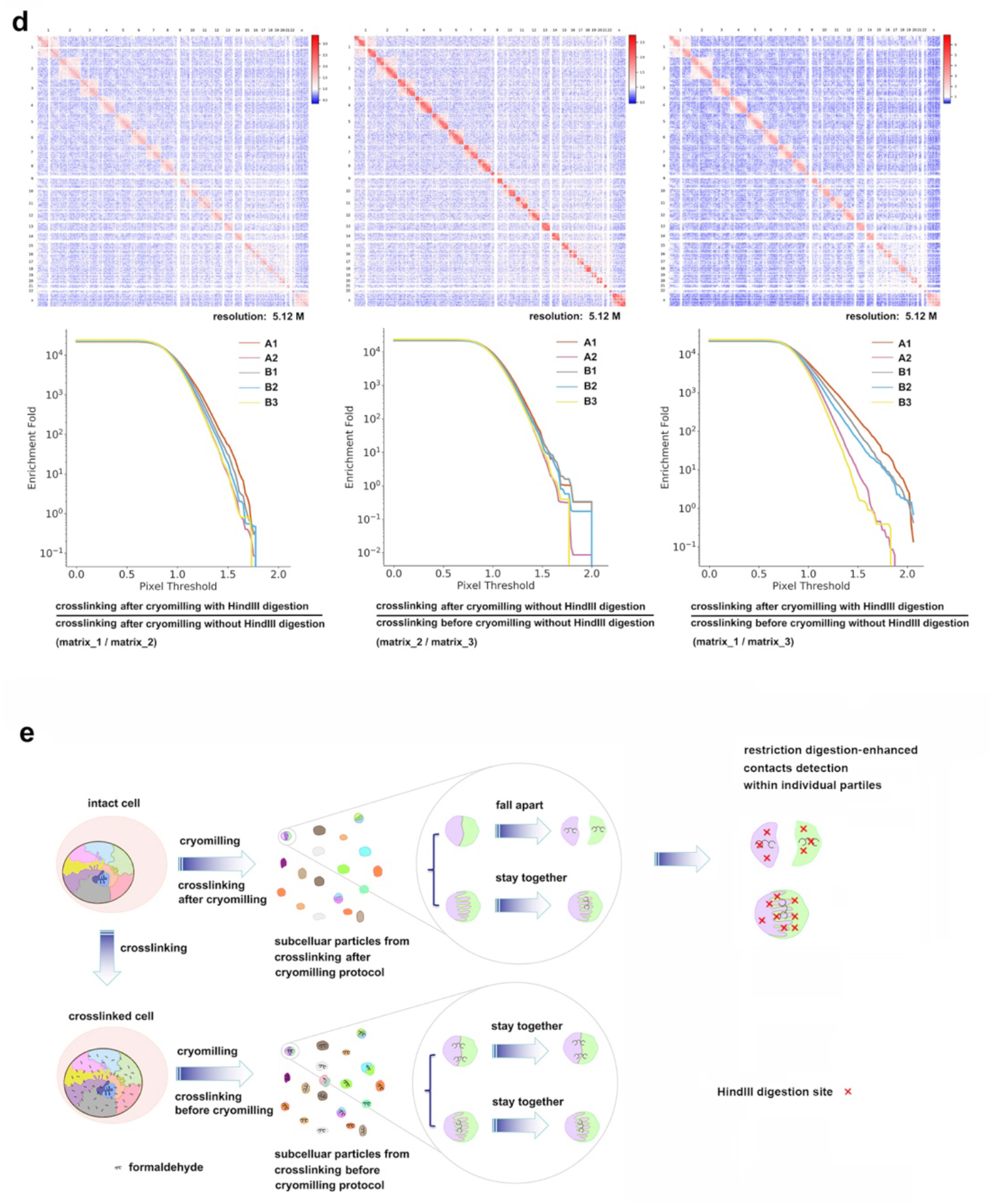
Effects of additional HindIII digestion on CTCC and the synergistic effect of “xlinking after cryomilling” and “HindIII digestion” on revealing stable contacts in CTCC experiments. **a**. Illustration of DNA breaks generated by cryomilling or HindIII digestion on cryomilled subcellular particles **b**. Same compartment pair frequency-pair distance curve comparison of CTCC without HindIII and CTCC with HindIII **c**. upper panel: Division matrices between crosslinking after cryomilling with HindIII digestion (matrix_1), crosslinking after cryomilling without HindIII digestion (matrix_2) and crosslinking before cryomilling without HindIII digestion (matrix_3). Lower panel: trans pixels’ subcompartment enrichment curve calculated from the division matrices mentioned above, respectively. **d.** Comparison of three sets of division matrix and corresponding subcompartment enrichment curve showing the synergistic effect of “xlinking after cryomilling” and “HindIII digestion” on revealing stable contacts **e**. Proposed principle of how crosslinking after cryomilling (crosslinking in vitro) and HindIII digestion enrich stable chromatin contacts in CTCC experiments.

**Supplementary Figure 7.**
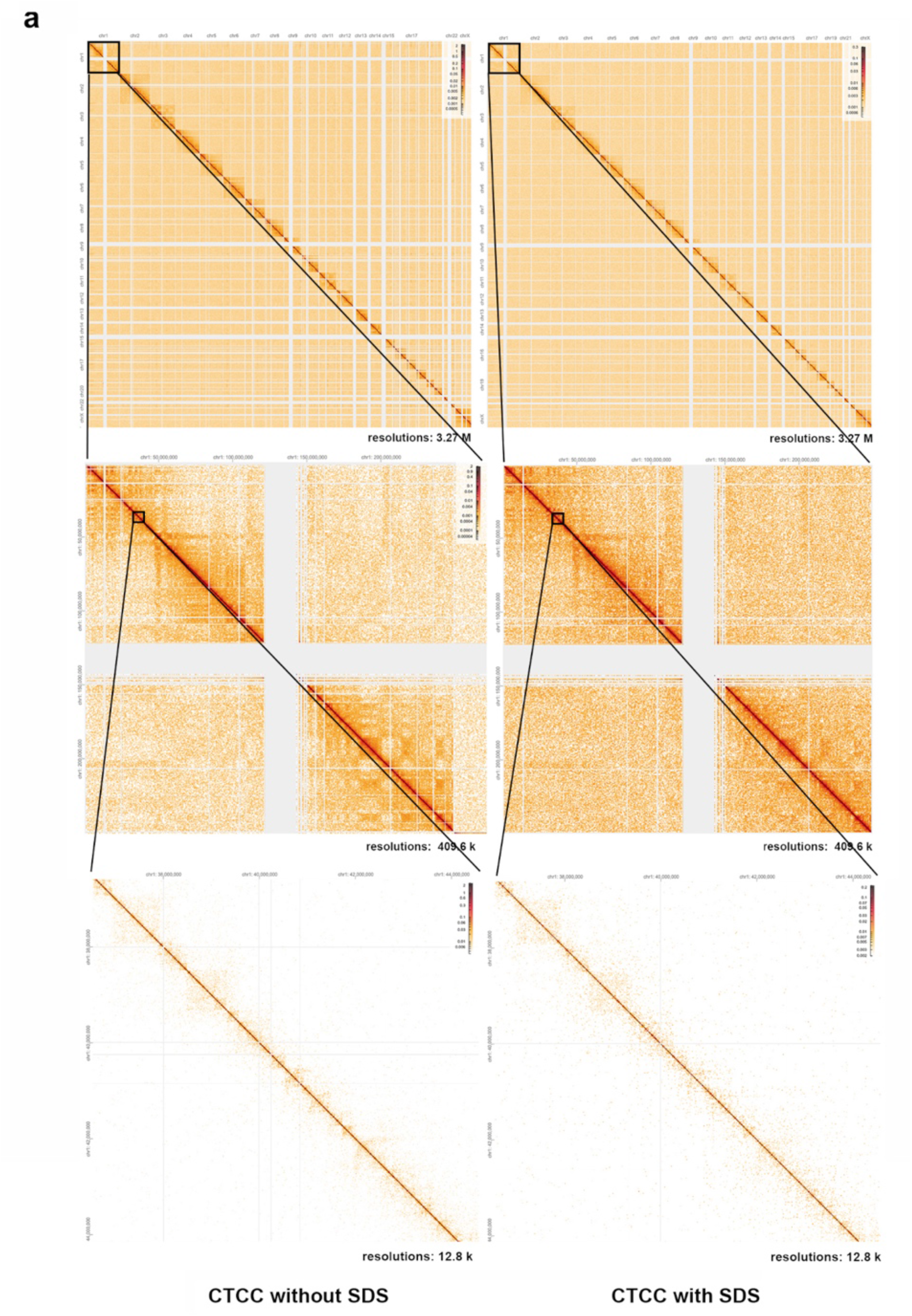
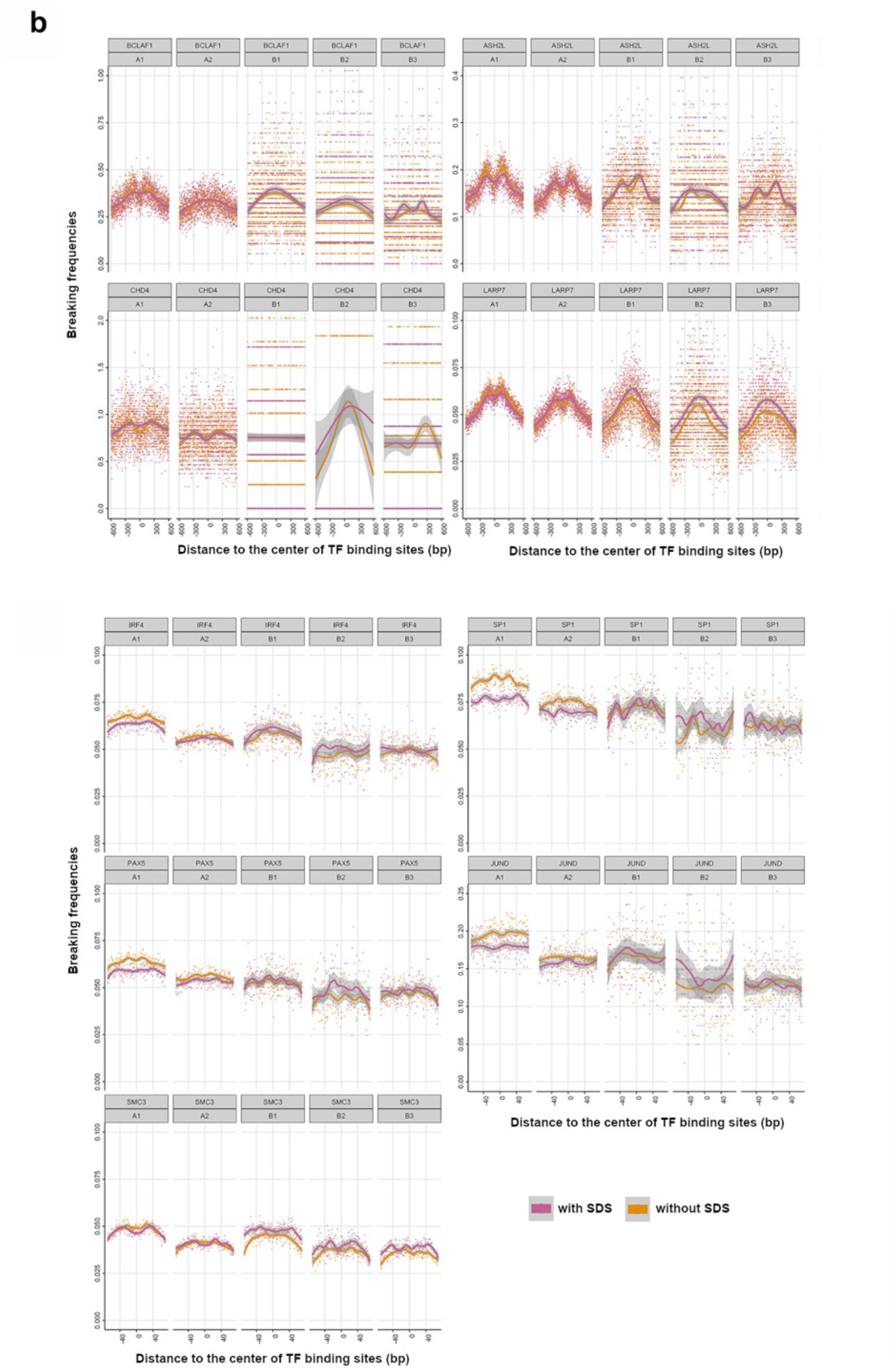
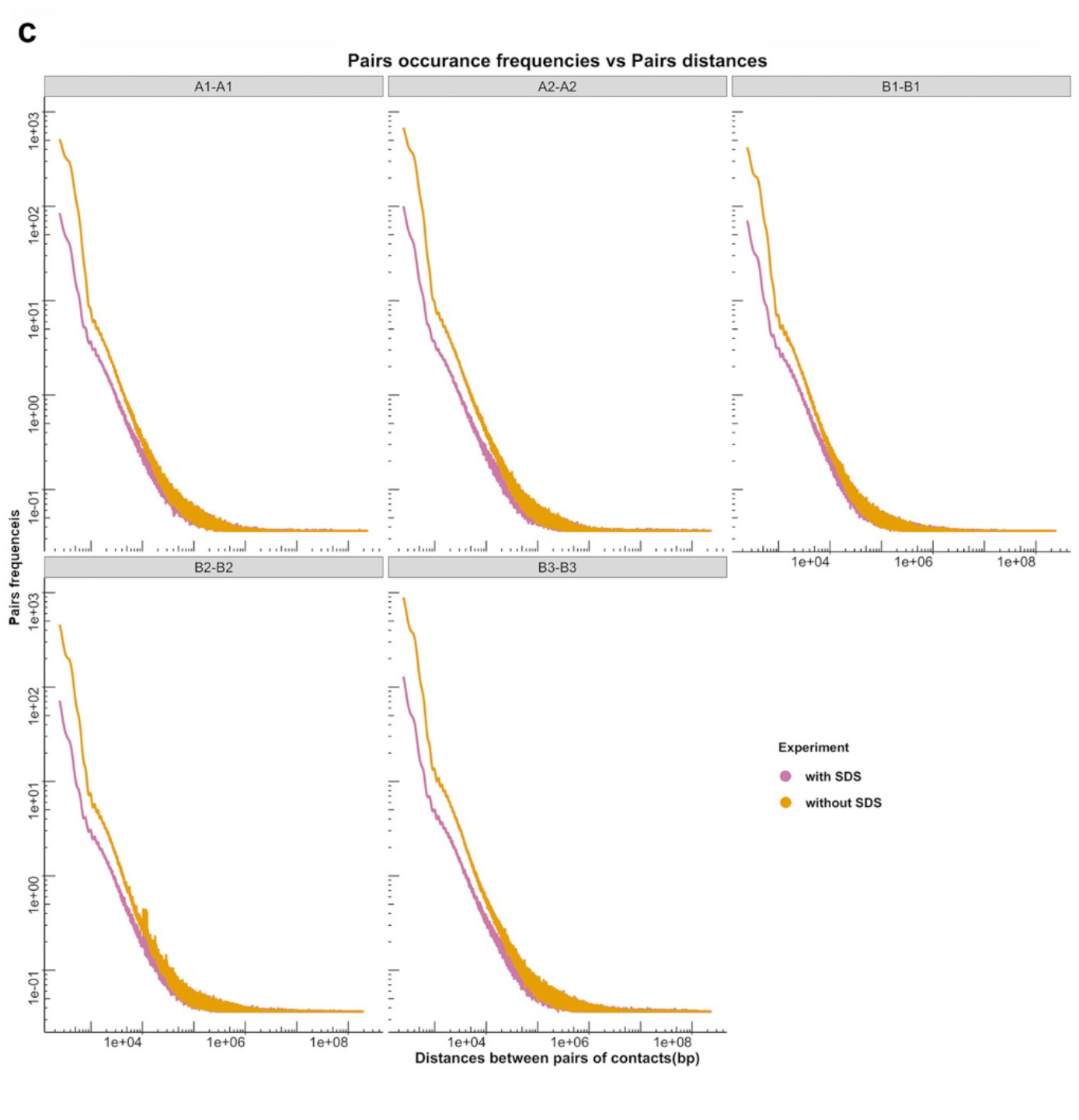
Effects of SDS treatment on CTCC experiments. **a**. Comparison of the contact matrices of CTCC experiment with and without SDS. **b**. The pileup distribution of mappable reads in CTCC experiment in terms of enrichment ratio of DNA breaks vs distance to transcription factors’ bindings sites. **c**. Comparison of Same compartment pair frequency-pair distance curve between CTCC with SDS and CTCC without SDS, both with crosslinking before cryomilling.

**Supplementary Figure 8.**
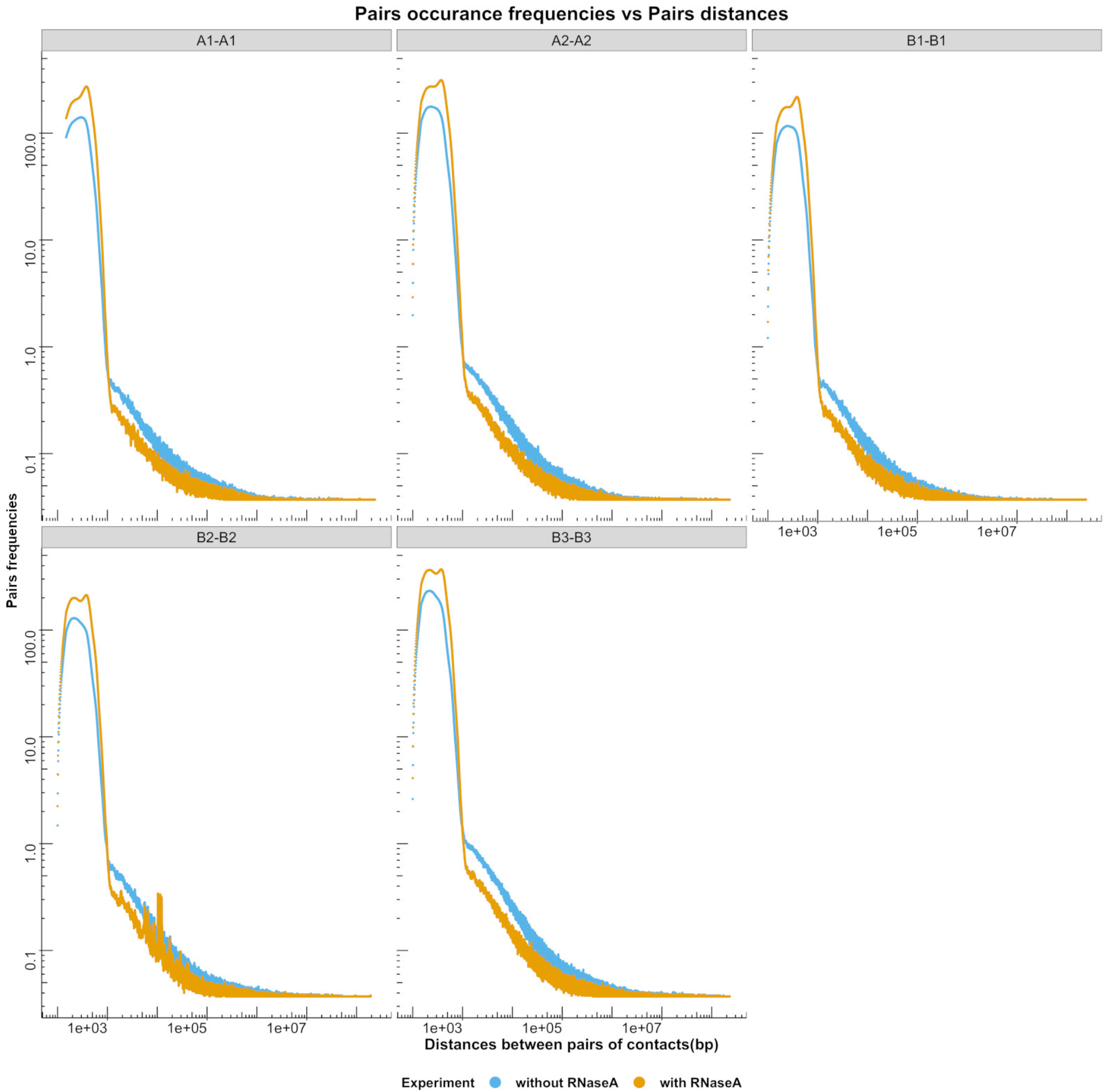
Effects of RNase A treatment on CTCC experiments. Comparison of same compartment pair frequency-pair distance curve between CTCC with RNase A and CTCC without RNase A, both with crosslinking after cryomilling (crosslinking in vitro).

**Supplementary Figure 9.**
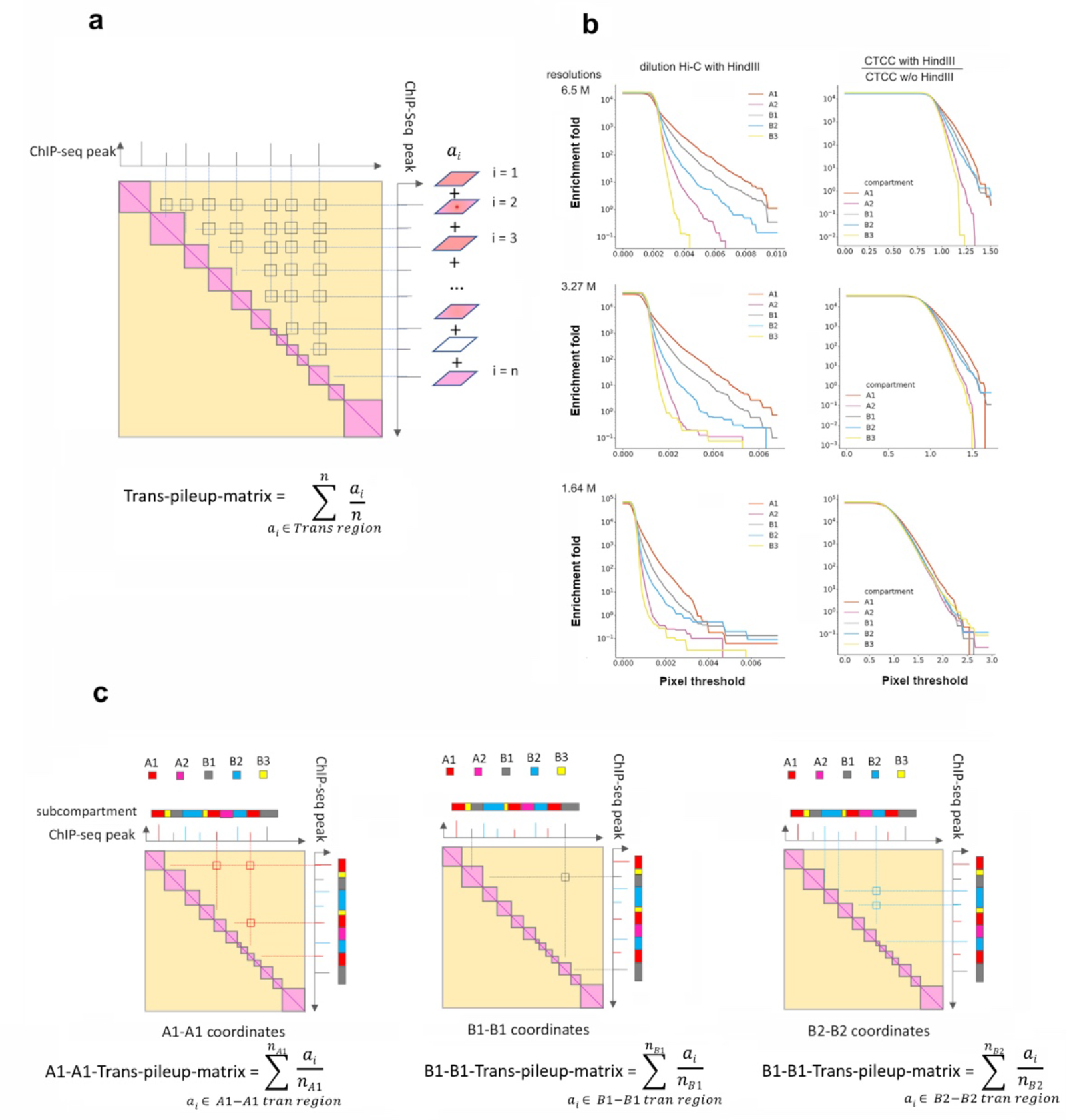
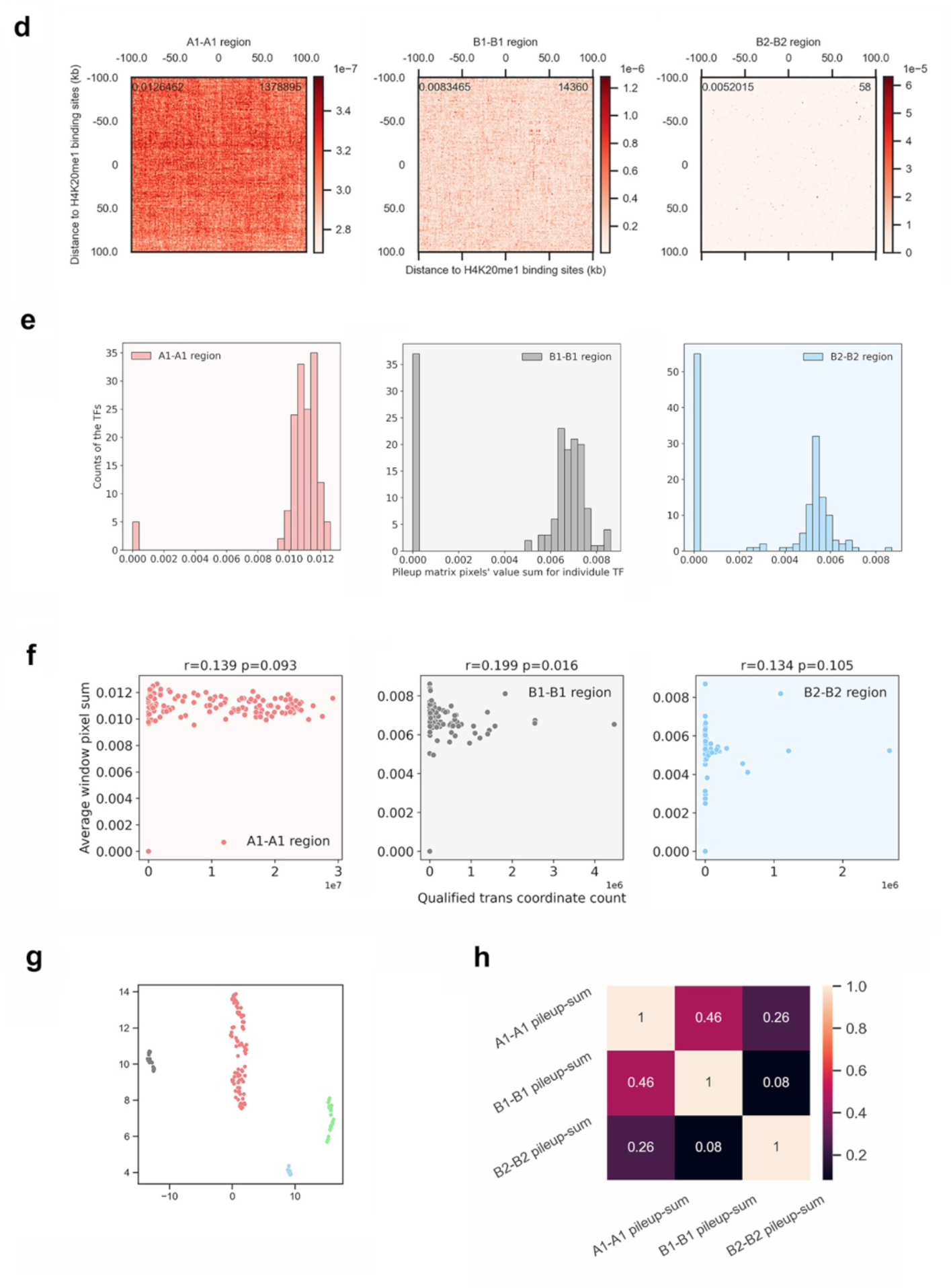
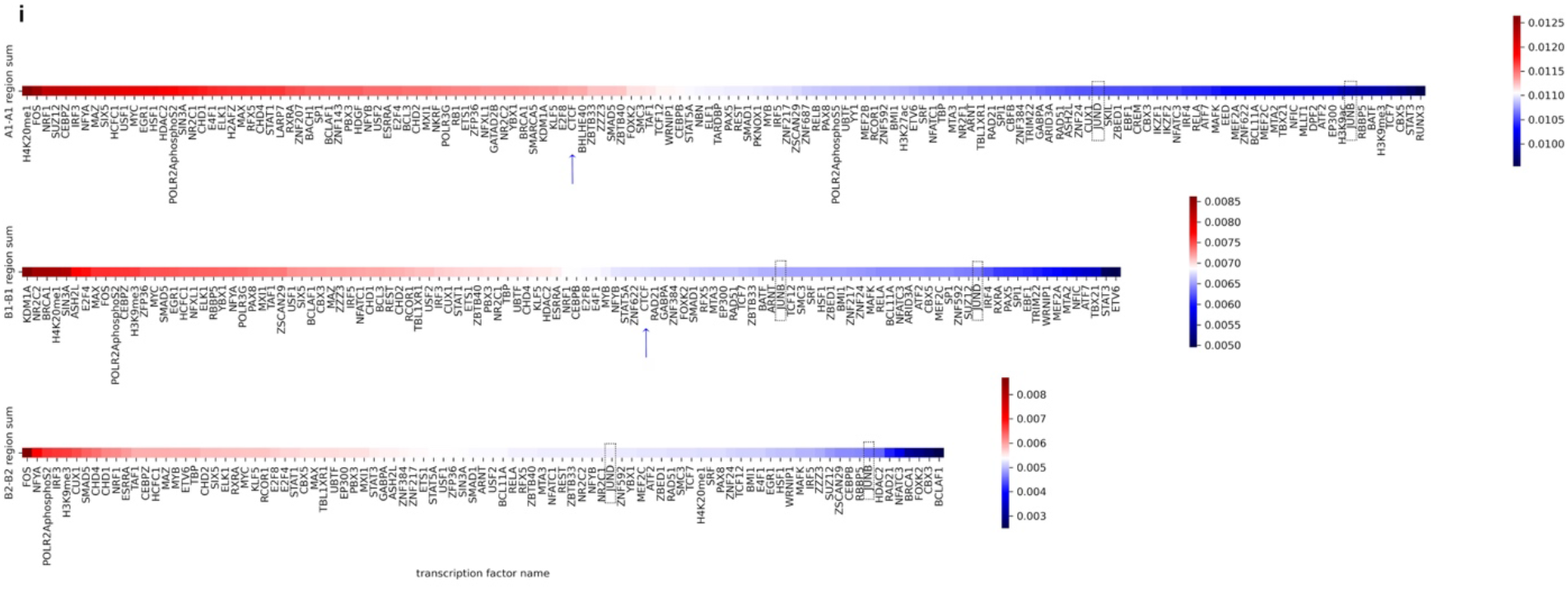
Trans pileup assay using 1kb resolution dilution Hi-C matrix file and known ChIP-Seq data for GM12878 cells. **a.** Algorithm illustration of trans matrix pileup according to ChIP-Seq peak position. **b.** Comparison of trans pixels’ subcompartment enrichment curve between dilution Hi-C (with HindIII) and division matrix of CTCC with HindIII CTCC without HindIII at different resolutions. **c**. Algorithm illustration of trans pileup matrix according to same subcompartment ChIP-Seq peak position. **d.** Example plot of trans pileup matrices according to the ChIP-Seq peak of H4K20me1 (numbers shown on the upper left, upper right are pileup-sum and total number of coordinates, respectively). **e**. Histogram showing the distribution of pileup-sum value of different nucleoproteins in terms of A1-A1, B1-B1 and B2-B2 pileup matrix. **f**. Scatter plot of pileup sum values vs total number of coordinates for individual nucleoproteins in terms of A1-A1, B1-B1 and B2-B2 pileup matrix. **g**. UMAP clustering of different nucleoproteins in terms of subcompartments bindings and pileup-sum. Grey: A1, B1 binder; light coral: A1, B1 and B2 binder; light blue: strict A1 binder with higher pileup-sum value; light green: strict A1 binder with lower pileup-sum value. **h**. Correlation matrix of pileup-sum values for A1-A1, B1-B1 and B2-B2 pileup matrices of individual nucleoproteins (only nucleoproteins bind all three subcompartments of A1, B1 and B2 are considered). **i**. Heatmap showing the ranking of nucleoproteins in terms of pileup-sum, with blue arrows highlighting the ranking position of CTCF, JUNB and JUND highlighted with dashed black boxes.

**Supplementary Video Comparison of buffer’s pH and ionic strength in maintaining the stability of the cell particles.** The same amount of cell particles from the “crosslinking before cryomilling” experimental protocol was stored in the same volume of PBS (10 mM Phosphate buffer, 150 mM NaC, pH 7.4, left tube) or 10 mM Tris-HCl, pH 8.0 (right tube) overnight at room temperature. Cell particles stored in PBS form visible large aggregates out of the suspension while those in 10 mM Tris-HCl remain homogenous.

## Materials and Methods

Cell growth, crosslinking and cryomilling Approximately 1.44 x 10^9^ GM12878 cells (Coriell Institute) grown in 1.8 liters of 1640 medium with 15% FBS were pellet down by centrifugation at 1000 rpm/20 min and washed with 1xPBS once. Cells were then resuspended at 50% (V/V) with isotonic 5% glucose solution and flash frozen by transferring with a Pasteur pipette into liquid nitrogen in a ‘drop to drop’ manner. The frozen droplets were then collected and stored at − 80 °C. For cells that require crosslinking before flash freezing, the 1xPBS washed cell pellets were resuspended in 45 ml of PBS into which formaldehyde solution was added to a final concentration of 1% and incubated with rotation at RT for 15 min. The crosslinking reaction was then quenched with addition of 2 M glycine solution to a final concentration of 170 mM and incubated further for 10 min/RT, followed by washing with PBS. Flashing freezing was then done as described above.

### Cryomilling of GM12878 cells

Cryomilling of frozen cells was performed under liquid nitrogen in a 50 ml milling jar using a Retsch PM100 planetary ball mill. To prevent heating of the sample the milling jar was placed inside a custom-made Polytetrafluoroethylene (PTFE) insulator jar during the whole milling process. The milling jar, PTFE insulation jar and all handling tools used for cryomilling were precooled in liquid nitrogen before use. 2 to 5 g of frozen cell beads were transferred into the precooled milling jar, and then by using forceps three precooled 20 mm diameter milling balls were placed in the jar. The jar was half filled with liquid nitrogen, the lid was placed on the jar, and the whole assembly was transferred to the planetary ball mill. One cycle of milling was performed with the following program: 450 rpm, 6 min grinding time, reverse-rotation each 45 sec and no interval break. After the cycle, the sample was cooled with liquid nitrogen and 20 mm milling balls were removed. Using forceps, 12 precooled 10 mm diameter milling balls were then placed into the milling jar. Ten cycles of milling were performed with the following program: 500 rpm, 8 min grinding time, reverse-rotation each 30 sec and no interval break. Between each cycle the milling jar along with the insulator was cooled with liquid N_2_ and packed cell powder was dislodged by a spatula. At the beginning of each milling cycle the milling jar with the cell powder was half filled with liquid nitrogen. At the completion of cryomilling, the 10 mm balls were removed, and ∼200 mg of cryomilled cell powder was aliquoted into 2 ml cryovials and stored at −80 °C until further usage.

### CTCC with in-vitro crosslinking (cryomilling before crosslinklinking)

For CTCC experiments starting from non-crosslinked cryomilled cell powder, approximately ∼200 mg of the cryomilled cell powder was thawed into 10 ml of 1 x PBS solution containing 1% formaldehyde, 1% Triton X^TM^-100 with vigorous shaking, and then incubated at RT for 15 min. 2 M Glycine solution was then added to a final concentration of 170 mM to quench the reaction. The cell particles were then pellet down by centrifugation at 2000 *g*/4°C for 15 min and were resuspended with 400 μl of 0.5% SDS and transferred to 1.5 ml Eppendorf tube and incubated at 65 °C for 10 min, with occasional pipetting to enhance solubilization of the cell particles. To attach biotin residues to the particles, 100 μl of freshly prepared 25 mM EZ-Link™ Iodoacetyl-PEG2-Biotin (IPB, Thermo Scientific) was added to the solubilized cell particles to a final concentration of 5 mM and incubated at RT for 30 min in the darkness. Free IPB was then quenched with addition of 2.5 μl of 2-mercaptoethanol to a final concentration of 60 mM and removed by exhaustively dialyzing against 20 mM Tris-HC, 50 mM NaCl, 0.5% SDS in a 3 ml Slide-A-Lyzer™ Dialysis Cassettes (Thermo Fisher Scientific) overnight. To prepare the particles for tethering to the magnetic beads, 10xCutSmart buffer (NEB), 20% Triton^TM^ X-100 solution and deionized water (dH_2_O) were added to the dialyzed cell particles suspension to contain a final concentration of 1xCutSmart, 1% Triton^TM^, 0.1% SDS, and incubated at RT for 10 min. 400 μl of Dynabeads™ MyOne™ Streptavidin T1 were washed with 1 ml of Wash-Buffer 1 (WB1) and was mixed with the cell particle suspension and incubated at RT for 30 min. For CTCC experiments with RNase A treatment, RNase A (Thermo Fisher) was added to 100 μg/ml to the cryomilled-cell-particles-T1-beads-mixture and incubated at RT for 30 min. The particles-tethered-magnetic-beads (hereafter referred to as ‘P_beads’) were then either digested with 1000 U of HindIII-HF (NEB) in 1 ml of WB1 at 37 °C with mild rotation for over 6 hrs, or not digested with HindIII for CTCC with cryomilling generated ends only, followed by incubating with WB1 supplemented with 5 mM biotin (Sigma-Aldrich) at RT for 15 min to block free biotin-binding sites on the beads and then washing with 1 ml of WB1 for 3 times. The P_beads were then incubated in 200 μl of Ends-Repair-Mix (ERM) to make blunt ends and then incubated with 1 ml of 0.1% SDS and washed twice with 1 ml of WB1. To add protruding A at the ends, the P_beads were incubated with 200 μl of Add-A-Mix (AAM) at 37 °C for 30 min and were then washed with 1 ml WB1 for 4 times. To ligate biotinylated-adaptors (hereafter referred to as Con2)(66), the P_beads were incubated with 200 μl of Adaptor-Ligation-Mix (ALM) and incubated at 18 °C for over 10 hrs and then washed with 1 ml of WB1 5 times. To phosphorylate the 5’ ends of the ligated adaptor, the beads were incubated with 100 μl of phosphorylation solution (PM) at 37 °C for 1 hr and was then mixed and incubated with 9 ml of ligation-mix (LM) at 18 °C to finish proximity ligation. Ligation product was then extracted as follows: the P_beads were resuspended in 500 μl of the DNA Extraction Mix (DEM) and incubated at 58 °C for more than 8 hrs, followed by decrosslinking with addition of 55 μl of 5 M NaCl and incubated at 68 °C for 2 hrs. The digested and de-crosslinked P_beads were then further extracted with 400 μl of Phenol-Chloroform-Isoamyl-Alcohol (PCI) once and 400 μl of Chloroform once. DNA was then precipitated with addition of 1/10 volume of 3 M sodium acetate (pH 5), 2 volumes of ethanol, incubated at −20 °C for 30 min, and then pellet down by centrifugation at 14800 *g*/4 °C for 15 min. The pellets were washed with 1 ml of cold 70% Ethanol twice, air dried and stored at − 80 °C.

### CTCC with in-situ crosslinking (crosslinking before cryomilling) with SDS

For crosslinking cells for cryomilling, 1.44 billion GM12878 cells from 1.8 liters of culture were pellet down by centrifugation at 1000 rpm for 15 min, washed once with 50 ml of 1xPBS and then resuspended in 50 ml of 1xPBS buffer containing 1% formaldehyde to crosslink the cells. Crosslinking was then quenched as described above. Crosslinked cells were then washed once with 50 ml 1xPBS and finally resuspended to 50%(V/V) with 5% glucose and flash frozen and cryomilled as described above.

For CTCC experiments using cryomilled cell powder from formaldehyde crosslinked cells, the experiments could be done with or without SDS solution treatment. For experiment with SDS treatment, ∼ 200 mg cryomilled cell powder from crosslinked cells was thawed into 1 ml of 1xPBS, pelleted by centrifugation at 2000 *g*/15 min and solubilized in 0.5% SDS at 62 °C for 10 min. P_beads were generated as described before. Ends-repair, A-addition, adaptor ligation and phosphorylation, and proximity ligation were carried out also as described before. Ligation product was then extracted, de-crosslinked as described above.

### CTCC with in-situ crosslinking (crosslinking before cryomilling) without SDS

For CTCC experiments without SDS treatment, ∼ 200 mg of the cryomilled cell powder from formaldehyde crosslinked cells was thawed into 1 ml of 10 mM sodium phosphate buffer, pH. 8.8 (PB), and dialyzed against 1 L of PB in a 3 ml Slide-A-Lyzer™ Dialysis Cassettes (Thermo Fisher Scientific) at RT for 3 hrs. To tether the cell particles to the magnetic beads, the dialyzed cell particles were mixed with 300 μl of Pierce™ NHS-Activated Magnetic Beads (Thermo Fisher), which was pre-washed with 1 mM hydrochloride solution, and incubated at RT for 2 hrs to generate the P_beads. To block unreacted sites, the P_beads were collected on a magnetic stand and resuspended in Wash-Buffer 2 (WB2) containing 10 mM Tris-HCl (pH 8.0), 0.5% Triton-X100 and incubated at RT overnight. The P_beads were then resuspended in 200 μl of ERM and incubated at RT for 1 hr followed by washing with 1 ml WB2 for 3 times and then 1 ml WB1 once. To add protruding A to the repaired 3’ ends, the P_beads were resuspended in 200 μl of AAM, and incubated at 37 °C for 1 hr, followed by washing with WB2 with 1mM EDTA three times and WB1 and 10 mM Tris-HCl (pH 8.0) once, respectively. To ligate Con2, the P_beads were resuspended in 200 μl of ALM and then incubated at 18 °C for over 10 hrs, followed by washing with 1 ml of WB2 with 1mM EDTA three times, 1 ml of WB1 once and 1 ml of 10 mM Tris-HCl (pH 8.0) once respectively. To phosphorylate the 5’ ends of the ligated adaptor, the P_beads were incubated with 100 μl of phosphorylation solution (PS) at 37 °C for 1 hr, and was then mixed and incubated with 9 ml of ligation-mix (LM) at 18 °C for over 10 hrs to finish proximity ligation. Ligation product was then extracted, de-crosslinked, pelletized and stored as described above.

### DNA purification and NGS library preparation

The DNA library for NGS sequencing was prepared as described before(66) with some revision. Briefly, the purified DNA pellets stored at −80 °C from the experiments were solubilized in 130 μl of 10 mM Tris-HCl, pH 8.0, 100 μg/ml RNase A (Thermo Fisher) at 37 °C for 40 min, transferred to microTUBE AFA Fiber Pre-Slit Snap-Cap (6×16mm) and sheared to around 400 bp with Covaris S2 (Intensity 4, duty cycle 10%, cycle per burst 200, treatment time 20 sx3). Sheared DNA was then mixed with 100 μl of 2 x Binding Buffer (2xBB) and 50 μl of T1 beads prewashed with 100 μl of 1 x Binding and Washing Buffer (1xBW). The mixture was then incubated at RT for 20 min and then washed three times with 600 μl of 1xBW, followed by washing with 600 μl of WB1 and 600 μl of 10 mM Tris-HCl, pH 8.0, respectively. The DNA-tethered T1 beads (hereafter referred to as D_beads) were then resuspended in 40 μl of ERM and incubated at RT for 30 min. The D_beads were washed by being resuspended with 1 ml of 1xBW, incubated at 55 °C for 2 min, which were repeated three times, and then washed once with 1 ml of WB1. To add protruding A to the 5’ end of the DNA, the D_beads were resuspended with 40 μl of AAM, incubated at 37 °C for one hr, followed by washing with 1 ml of 1xBW three times with heating to 55 °C as described above, and then 1 ml of WB1 once and 1 ml of 10 mM Tris-HCl, pH 8.0 once, respectively. To ligate the Y-adaptor, the D_beads were resuspended in 30 μl of Y_adaptor_Ligation_Mix (YLM) and incubated at 18 °C for 16 hrs, followed by washing with 1 x BW and heated to 55 °C three times as described above and then washing with 1 ml of 10 mM Tris-HCl, pH 8.0, and finally resuspended in 50 μl of 10 mM Tris-HCl, pH 8.0. The D_beads suspension could then be stored at −80°C for further usage. To amplify the library, 5 μl of the D_beads suspension was mixed with 2.5 μl of forward primer SeqPrimer_F, 2.5 μl of barcoded Index primer, 15 μl of dH_2_O, and 25 μl of NEBNext Ultra II Q5 Master Mix (New England Biolabs), and incubated on a thermocycler at 98°C/30s, (98°C/10s, 65°C/75s) for 8∼15 cycles, and 65°C/5 min. The amplified product was size-selected with SPRIselect beads (Beckman Coulter) to select the dsDNA between 300∼500 bp for NGS. NGS experiments with paired-end- 150bp (PE150) reads were done at the Technology Center for Genomics & Bioinformatics (TCGB), University of California, Los Angeles.

### Buffers and Premixes

Wash Buffer 1 (WB1): 0.5% Triton X-100, 1xNEBuffer 2

Wash Buffer 2 (WB2): 0.5% Triton X-100, 10 mM Tris-HCl, pH 8.0

End Repair Mix (ERM): 1xT4 ligase buffer with 1 mM ATP, 0.5 mM dNTP, 0.5 U/μl T4 PNK, 0.12 U/μl T4 DNA Polymerase, 0.5 U/μl Klenow-Fragment, 1% Triton X-100 and 1 mM Biotin (Sigma-Aldrich)

A Addition Mix (AAM): 1xNEBuffer2, 0.5 mM dATP, 1 % Triton X-100, 0.375 U/μl Klenow-Fragment (3’ → 5’ exo-)

Adaptor Ligation Mix (ALM): 9 μM adaptor, 1xT4 ligase buffer with 1 mM ATP, 0.5% Triton X-100, 2.5 U/ μl T4 DNA Ligase (NEB), 1 mM Biotin

Phosphorylation Mix (PM): 1xT4 ligase buffer with 1 mM ATP, 0.5% Triton X-100, 1 U/μl T4 PNK (NEB)

Ligation Mix (LM): 1xT4 ligase buffer, 0.5% Triton X-100, 2 U/μl T4 DNA Ligase (NEB)

DNA Extraction Mix (DEM): 1xDNA Extraction Buffer (1xDEB), 1 mg/ml Proteinase K (NEB) and 1 mM Biotin

YLM: 1xT4 ligase buffer with 1 mM ATP, 1 μM Y adaptor, and 33 U/μl T4 DNA ligase

## Data analysis

### CTCC data processing and visualization

CTCC data processing and contact matrix visualization For Hi-C data analyses, sequencing data were splitted into blocks containing 2 million reads each, and then mapped to the reference genome with Bowtie2(67), processed to Hi-C pairs using the HiC-Pro pipeline(68). Pooled pairs files were randomly trimmed to the desired pair number with the python code “data_trimming.py” and then loaded to the Hi-C matrix with the cooler package(69). The cooler files generated were then zoomified and balanced using the iterative correction and eigenvector decomposition (ICE) method(70) with the cooler package to generate the multi-resolution ‘mcool’ file and loaded to HiGlass(71) to visualize the contact map. Division matrix were either generated in HiGlass or using the python code “cool_over_cool.py”

### Genomic DNA breaking position analyses

For DNA breaking position analyses of the CTCC or Hi-C experiments, sequencing reads containing the ligation junction sequence ‘AGCTGAGGGATCCCTCAGCT’ (for CTCC) or ‘AAGCTAGCTT’ (for dilution Hi-C with HindIII), or ‘GATCGATC’ (for dilution Hi-C with MboI) were trimmed off the ligation junction sequence either from 3’ or 5’ and saved to a new fastq file with Cutadapt(72). The trimmed sequencing reads in the newly generated fastq file were then mapped to the reference genome with bwa mem(73) to generate the bam file, from which the breaking positions were read extracted using pysam(74) with the following algorithm: breaking position is the mapping position, if the trimmed sequencing read is mapped to the forward strand and the ligation junction lies at the 5’ end of trimmed reads, or if the sequencing read is mapped to the reverse strand and ligation junction lies at the 3’ of the trimmed reads. Breaking position is the mapping position plus the aligned read’s length, if the sequencing read is mapped to the forward strand and the ligation junction lies at the 3’ end of trimmed reads, or if the sequencing read is mapped to the reverse strand and ligation junction lies at the 5’ of the trimmed reads. For the calculated breaking positions, the corresponding sub-compartment that the breaking position belongs to was obtained by querying a tabix-indexed(75) bed file containing the subcompartment information of the GM12878 cell(6). The above algorithms were realized with the python code “cal_breaking_hic.py”.

For MNase-seq and SPRITE data, the DNA breaking position was obtained directly from the mapped bam file. The breaking position is the mapping position if the read is mapped to the forward strand, or the mapping position plus the mapped read length, if the read is mapped to the reverse strand. The DNA breaking positions and corresponding subcompartments information were obtained with the python code “breaking_subcomp_tagging.py”. For DNA breaking density distribution analysis, the breaking position file was indexed with tabix and then queried with the python code “breaking_RPKM.py” with designated bin size. For visualization of the DNA breaking position distribution in the genome browser, the breaking positions were aggregated and then converted to a bigwig file, which was then loaded to the IGV genome browser installed on a local computer.

### Raw pairs’ “Signal” and “Noise” analysis for CTCC and Hi-C data

CTCC and Hi-C’s signal and noise were compared in terms of overall same subcompartment pair profile and in terms of each genomic regions (Hi-C matrix bins) contact profiles.

For pairs’ overall “signal” and “noise” analysis, “Cis Signal”, “Cis Noise”, “Trans Signal”, and “Trans Noise” for all 5 known major subcompartments, A1, A2, B1, B2 and B3 were calculated as follows: Pairs with read_1 and read_2 belong to the same subcompartment and the same chromosome were counted as “Cis Signal” for that particular subcompartment. Pairs with read_1 belong to a particular subcompartment and read_2 belong to a different subcompartment and the same chromosome were counted as the “Cis Noise” for that particular subcompartment. Pairs with read_1 and read_2 belong to the same subcompartment but different chromosomes were counted as “Trans Signal” for that particular subcompartment. Pairs with read_1 belong to a particular subcompartment and read_2 belong to a different subcompartment and a different chromosome were counted as the “Trans Noise” for that particular subcompartment.

For analysis of the contact’s “Signal” and “Noise” for each genomic region (Hi-C matrix bin), “bin’s_cis_same_compartment_pair_ratio” and “bin’s_trans_same_compartment_pair_ratio” were calculated as follows:

For each bin of the horizontal (or vertical) axis of the contact matrix: contact pairs with read_1 fall into that bin and read_2 fall into the same chromosome were summed to calculate the “bin’s_total_cis_pair_counts” and contact pairs with read_1 fall into that bin and read_2 fall into another chromosome were summed to calculate the “bin’s_total_trans_pair_counts”. Contact pairs with read_1 fall into that bin and read_2 that is the same subcompartment and mapped to the same chromosome as read_1 was counted as “bin’s_cis_same_compartment_pair_counts”. Contact pairs with read_1 fall into that bin and read_2 that is the same subcompartment but mapped to a different chromosome as read_1 was counted as “bin’s_trans_same_compartment_pair_counts”. For each bin, “bin’s_cis_same_compartment_pair_ratio” and “bin’s_trans_same_compartment_pair_ratio” were then calculated using the following equation:

”bin’s_cis_same_compartment_pair_ratio” = “bin’s_cis_same_compartment_pair_counts”/ “bin’s_total_cis_pair_counts”.

”bin’s_trans_same_compartment_pair_ratio” = “bin’s_trans_same_compartment_pair_counts”/ “bin’s_total_trans_pair_counts”

### 1D-pileup assay of DNA breaking positions to reference sites

For pileup relative to nucleosome centers, the following processes were taken: First, the bigwig file containing the information of nucleosome positions(31) were converted to bedgraph format using ‘bigWigToBedGraph’(76), from which the peak center was obtained with the python code “cal_bw_peak_pos.py” and the breaking position relative to the nearest center of the nucleosome was calculated with the python code “calc_breaking_nc_dist.py”. Second, the aggregated breaking positions relative to the center of the nucleosome were calculated with the python code “aggr_breaking_nc_dist.py”. For the 1D-pileup assay of distribution of breaking position relative to transcription factor (TF) binding sites, TF binding center was obtained with ChIP_seq’s narrow bed file from the encode database(49, 50), and the pileups of genomic DNA breakings relative to the TF binding center were calculated with the python code “calc_breaking_TF_dist.py” and visualized with ggplot2(77).

### Pair statistics and pair frequency-pair distance analysis

Pairs statistics analysis was done either with the HiC-Pro pipeline, or with the python code “pairs_subcomp_tagging_rev1.py” and pair-frequency-pair-distance curve and same-compartment-pair-frequency-pair-distance data was generated with “cal_pair_dist_freq.py” and visualized with ggplot2.

### Trans pixels’ subcompcompartment enrichment curve calculation and plotting

To calculate trans’ pixels’ subcompartment enrichment fold, each trans pixel was parsed for the subcompartment contents it covers by averaging the covered horizontal and vertical subcompartment length and normalized by the total subcompartment length (**Supplementary Figure 6c**). For plotting the trans pixels’ subcompcompartment enrichment curve, pixel threshold values were calculated by making 50 equal spacings from the lowest value in the trans matrix to the highest value in the trans matrix, which were devoid of ‘numpy.nan’ and ‘numpy.inf’. At each threshold, only trans pixels with value exceeding the threshold were included for calculation of enrichment fold for a particular subcompartment, which was calculated by summing up all pixels’ enrichment fold value for that subcompartment.

### Matrix division and pileup

Division matrix was either generated and visualized within the HiGlass browser or using the python code “cool_over_cool.py”. For matrix division, cool files from raw pairs file with the same “valid pairs counts” were used, which was generated by the python code “data_trimming.py” that can randomly trim the “. allValidPairs” file to the desired pair counts.

For TAD(4) boundary detection, insulation score was calculated using cooltools (https://github.com/open2c/cooltools) and insulation valley positions were calculated with the python code “cal_bw_peak_pos.py”. Cis 2-D pileups at the TAD boundaries and know loop positions(6) of the contact matrix were done using “Coolpup.py”(78). For trans pileup according to known ChIP-seq peaks for the GM12878 cell line, high resolution in-situ Hi-C map of GM12878 (4DNFIXP4QG5B.mcool from Lieberman Aiden lab) at 1 kb resolution was used in combination with the narrow bed files from the Encode portal(49, 50). “Transpileup_coord_generator.py” was used to generate the coordinate according to the following rules. Each narrow bed file was filtered for qualified peaks with FDR (q value) less than 0.05. To simplify the algorithm for coordinate generation, only one ChIP-seq file was considered at a time (e.g., Fos-Fos, H4K20me1-H4K20me1, but not Fos-H4K20me1). Only peaks belonging to the same subcompartment were paired to generate the submatrix center coordinates (**Supplementary Figure 9c**), which define potential submatrices that were further screened for those that only contain the trans interactions. Even with such criteria, pileups of TF peaks’ ‘crossings’ in trans proves very challenging because of the huge number of coordinates and the limited speed of the process and was achieved with splitting the coordinate file to a bunch of smaller files and multiprocessing. Python code “matrix_pileup.py” was used to do the pileup that qualified matrices specified with the coordinates with a pad value of 201 kb were added up and averaged by dividing the total number of the matrices.

## Contributions

L.C., F.A., J.A. and M.R. conceived the idea; J.X., S.K., and L.C. designed the experiment; J.X., Y.K. and X.L. did the cell culturing; S.K. did the cryomilling; J.X. did the CTCC experiments, and did the data analysis together with N.H.

## Data and computer code availability

The raw sequencing data and processed pairs, matrix files were deposited to the 4DN Data Portal with the dataset name ‘cryomilling TCC protocol variations’, with the title of 4DNESUGKCFC5, 4DNESHTTKZYB, 4DNESXF7RIFP, 4DNESXLN6HC6, 4DNES8MJ5EYG, 4DNESDG6WDIU and 4DNESNHBSICL, respectively. The python codes used were deposited to the GitHub page https://github.com/JiangXu123/CTCC_Related_Analysis_Tools.

## Acknowledgement

Data analyses were done at the Center for High-Performance Computing of University of Southern California. We would like to thank Yang Zhang (Jian Ma Lab at Carnegie Mellon University) for suggestions on data analysis, Nichola Servant (Institut Curie) and Luigi Manna (Department of Biological Sciences, University of Southern California) for helping to set up the HiC-Pro Pipeline, Nezar Abdennur (Leonid Mirny Lab at MIT) for helping with cooler related data analysis and Peter Kerpedjiev (Nils Gehlenborg Lab at Harvard Medical School) for Hi-Glass related data visualization.

